# Apparent simplicity and emergent robustness in bacterial cell cycle control

**DOI:** 10.1101/2023.01.16.524295

**Authors:** Sander K. Govers, Manuel Campos, Bhavyaa Tyagi, Géraldine Laloux, Christine Jacobs-Wagner

## Abstract

To examine how bacteria achieve robust cell proliferation across diverse conditions, we developed a method that quantifies 77 cell morphological, cell cycle and growth phenotypes of a fluorescently-labeled *Escherichia coli* strain and >800 gene deletion derivatives under multiple nutrient conditions. This approach revealed extensive phenotypic plasticity and deviating mutant phenotypes were often found to be nutrient-dependent. From this broad phenotypic landscape emerged simple and robust unifying rules (laws) that connect DNA replication initiation, nucleoid segregation, FtsZ-ring formation, and cell constriction to specific aspects of cell size (volume, length, or added length). Furthermore, completion of cell division followed the initiation of cell constriction after a constant time delay across strains and nutrient conditions, identifying cell constriction as a key control point for cell size determination. Our work provides a systems-level understanding of the design principles by which *E. coli* integrates cell cycle processes and growth rate with cell size to achieve its robust proliferative capability.

## Introduction

Bacteria are notorious for their proliferative capabilities, which have direct implications in medicine, agriculture, and industry. Like all cells, bacteria must grow in size, replicate their DNA, segregate their genome, and divide in order to multiply. These processes must be coordinated to ensure faithful self-replication across all cells^1^. Yet most bacteria, including the best-studied model *Escherichia coli*, seem to lack many of the cell cycle checkpoints and regulatory systems (e.g., cyclins, cyclin-dependent kinases, mTOR) that ensure smooth cell cycle progression in eukaryotic cells^2–4^. Despite this, bacterial replication is remarkably faithful (i.e., virtually error-free), with each mother cell producing daughter cells that are themselves competent for self-replication. For many bacteria, cellular replication is also robust to environmental fluctuations. For instance, *E. coli* and other bacteria can multiply under a vast array of nutritional conditions, exhibiting large variations in doubling times (< 20 min to days) and cell volume (up to five-fold for a single species)^5–7^. In fact, under nutrient-rich conditions, *E. coli* achieves the feat of dividing faster than it can replicate the genome by initiating DNA replication in one or two preceding division cycles^8^. Even then, cellular replication remains faithful.

How bacterial cells control cell cycle progression across diverse conditions is not well understood. In this study, we hypothesized that bacteria like *E. coli* are so adaptable because, at the systems level (i.e., on average), cells follow the same simple rules of cell cycle control across nutrient conditions and other perturbations. We propose to call such unifying rules “cell cycle laws”, by analogy with the growth laws that have been proposed previously. A classic example is the nutrient-imposed growth law, which describes that the average cell volume (V) of a bacterial population scales exponentially with the average growth rate (α) across nutrient-rich conditions (V ∝ e^α^)^5^A To our knowledge, the only known cell cycle law in bacteria applies to the initiation of DNA replication, which was first proposed in the 1960s^9^. It stipulates that in *E. coli*, DNA replication initiates at a constant volume (V_i_) per chromosomal origin of replication (*ori*). This cell cycle law was deduced from the nutrient-imposed growth law under the assumption that the C+D period (time of DNA replication + time between the end of DNA replication and cell division) is constant^9^. However, this assumption only holds under the nutrient-rich (fast-growth) regime^8^. To account for the scaling of the C+D period with the generation time in nutrient-poor (slow-growth) conditions^10–14^, the nutrient-imposed growth law has recently been expanded to include a variable C+D period (V ∝ e^α*(C+D)^ = 2 ^(C+D)/τ^; with τ being the average doubling time)^14,15^. This expanded relationship, referred to as the general growth law^14^ hereafter, relates the average cell volume of a population to its average number of *ori* (given by 2 ^(C+D)/τ^). The relation indicates a constant V_i_/*ori*, as V = V_0_ * 2 ^(C+D)/τ^ (with V_0_, the average cell size per *ori* or the unit cell volume, being directly proportional to V_i_/*ori*; V_0_ = V_i_/*ori* * ln(2))^14,15^. However, this constancy of V_i_/*ori* remains under debate as some studies support it^13–18^ while others do not^19–24^. This has also led to the proposal of an alternative relationship in which cell volume scales linearly with α * (C + D)^21^.

While the interplay between DNA replication initiation and cell size or growth rate has received much attention, it remains to be seen whether other cell cycle events follow unifying rules that reflect dependencies on cell size or growth rate. Below, we describe a large-scale phenotyping approach that allowed us to simultaneously extract quantitative features related to cell size, cell morphology and cell cycle progression across a wide array of perturbations in *E. coli*. This high-content cell data combined with growth rate measurements enabled us to map the phenotypic space of cell morphogenesis, population growth and cell cycle functions in *E. coli*. In addition to testing previously proposed relationships, our analysis suggests four new cell cycle laws. Taken together, our study provides population-level understanding of how *E. coli* robustly regulates and integrates cell cycle progression and growth rate through cell size-related design principles.

## Results

### Quantification of *E. coli* cell morphology and cell cycle events across nutrient conditions and large-scale genetic perturbations

To observe the interplay between cell morphology and different cell cycle events in *E. coli*, we used a strain harboring two chromosomally-encoded fluorescent fusions: one to the divisome component FtsZ^25^ and the other to the DNA replication marker SeqA^26^ (Figure 1A). We grew this strain under 41 different carbon source conditions (Table S1, Figure 1B, and S1A), giving rise to cultures of different average cell sizes (from ~1 to ~5 μm^3^) and doubling times (from ~20 min to ~3 h). For each condition, we stained exponentially growing cells with the DNA dye DAPI to visualize the nucleoids. We developed a pipeline for quantitative image analysis to extract singlecell features and calculate population averages for all extracted features from phase contrast and fluorescence microscopy snapshots (Figure 1A, see Methods). To determine the relative timings of each cell cycle event, we used an established method^19,27–29^, referred to here as the relative timing inference (RTI) method, which is based on the fraction of cells that have passed a specific cell cycle stage (Figure 1A). Several laboratories^30–33^, including ours (see Methods), have verified this method through comparison with time-lapse experiments. We used this RTI approach in combination with the fluorescent markers (FtsZ-Venus^SW^, SeqA-mCherry, and DAPI-stained nucleoids) to extract the relative timings of six cell cycle events: initiation and termination of DNA replication, FtsZ ring formation, initiation of cell constriction, initiation of nucleoid constriction, and nucleoid separation (Figure S1B-C, see Methods). In parallel, we obtained growth rate measurements from population growth curves (Figure 1A, see Methods), which were then used to transform relative cell cycle timings into absolute times.

**Figure 1.**
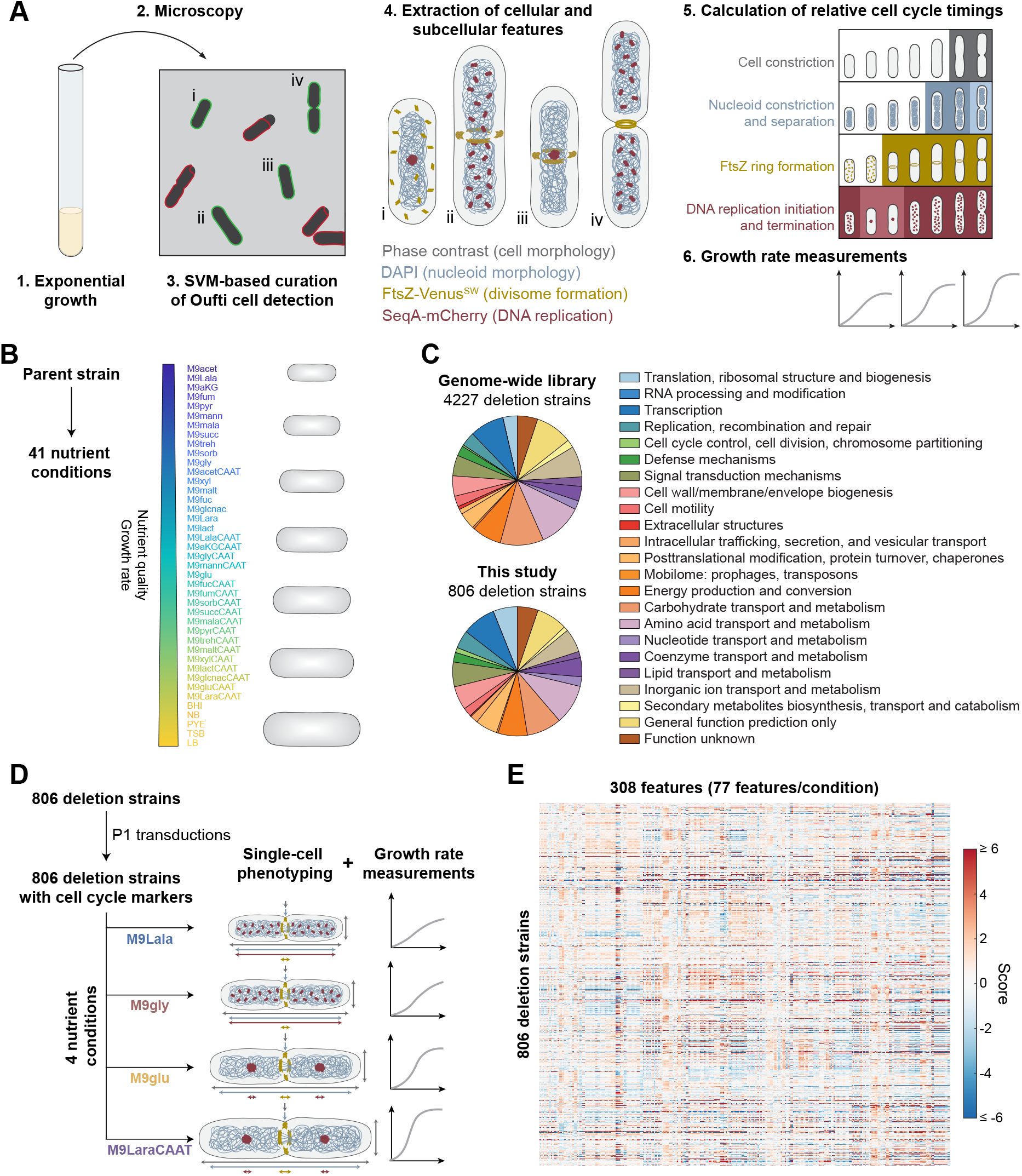
Experimental pipeline. A. Schematic showing the experimental pipeline, including the different steps of the microscopy-based phenotyping approach and the population growth measurements. B. Schematic showing the 41 nutrient conditions (Table S1) in which the parent strain was grown. The media differ in carbon quality and give rise to different growth rates and cell sizes. C. Pie charts representing the relative distribution of clusters of orthologous groups (COGs) categories among the gene deletion strains of the library used in this study versus those in the genome-wide Keio collection. D. Schematic showing the experimental approach for the generation and use of our gene deletion library, grown in four different nutrient conditions. E. Heatmap showing the feature scores (*s*) of the 806 deletion strains across the four nutrient conditions. See also Figures S1 and S2.

Modulating cell size by altering the nutrient composition of the growth medium is a common approach, but it has two limitations. First, regardless of the number of different media used, they all affect metabolism and, as such, they only represent one type of perturbation. Second, changes in nutrient quality result in correlated effects between growth rate and cell size^5,6^, preventing their uncoupling. To address these limitations and expand the phenotypic space, we used information from our previous genome-wide microscopy study of the *E. coli* Keio collection^34^ to select 806 gene deletions associated with one or more defects in cell/nucleoid morphogenesis and/or growth in M9 medium containing glucose, casamino acids, and thiamine (M9gluCAAT) (see Methods). As before^34^, we verified the reproducibility of imaging and measurements (e.g., cell length and width) across these 806 gene deletions in M9 medium containing glycerol (M9gly) (Figure S1D, see also Methods). Then, we individually transduced the gene deletions into our parent strain. Our subset of deletion strains was as diverse as the complete genome-wide library in terms of the distribution and number of different cellular processes that are impacted by the gene deletions (Figure 1C). Gene deletions that affect different processes effectively functioned as independent perturbations. This large number of independent and diverse perturbations enabled us to average out any specific effect conferred by a given mutation and to confidently identify dependencies between the different features of the system^34–37^.

To further increase the sampling of the phenotypic diversity available to *E. coli*, we analyzed our mutant library in four different nutrient conditions (Figure 1D): an M9 buffer with L-alanine (M9Lala), glycerol (M9gly), glucose (M9glu), or L-arabinose with casamino acids and thiamine (M9LaraCAAT). The nutrient conditions varied substantially in carbon source quality, giving rise to different average growth rates, cell sizes, and DNA replication regimes (discrete vs. overlapping DNA replication cycles)^5,6,8^. In total, we imaged approximately 4,300,000 individual cells, yielding 1384 ± 380 cells/strain/nutrient condition. Quantification of cellular features and cell cycle timings revealed that this mutant library generated broad phenotypic diversity within each nutrient and, thus, metabolic condition, as illustrated with key features in Figure S2A-B. For comparison, we imaged ≥ 115 replicates of the parent strain under the same four nutrient conditions (Figure S2A).

In total, we measured 77 different average features for each deletion strain in each nutrient condition (Figure 1E). We also calculated a normalized score (*s*) for each feature of a strain in each nutrient condition (see Methods). This score represents the extent to which a mutant strain deviates from the parent replicates in terms of standard deviations (SDs) away from the median.

### Cell length and width do not co-vary across large-scale perturbations

We first examined how cells control their size and shape. This question has been the subject of many studies in *E. coli* and other microbes. However, these investigations have led to conflicting conclusions about which dimensions of cell morphology (cell length, volume, surface area, the surface-area-to-volume ratio, or aspect ratio) cells keep constant^38–47^. These studies employed either a small number of perturbations (genetic, chemical or metabolic) or specific perturbations (targeting only a single protein or a small subset of proteins). Therefore, we used our large dataset of measurements to revisit this biological question by examining the relationship between cell length and width across perturbations. A constant aspect ratio predicts a positive relationship between cell length and width, whereas we expect different negative relationships for a constant cell volume, surface area, or surface-area-to-volume ratio, as depicted by the dashed lines in Figure 2A.

**Figure 2.**
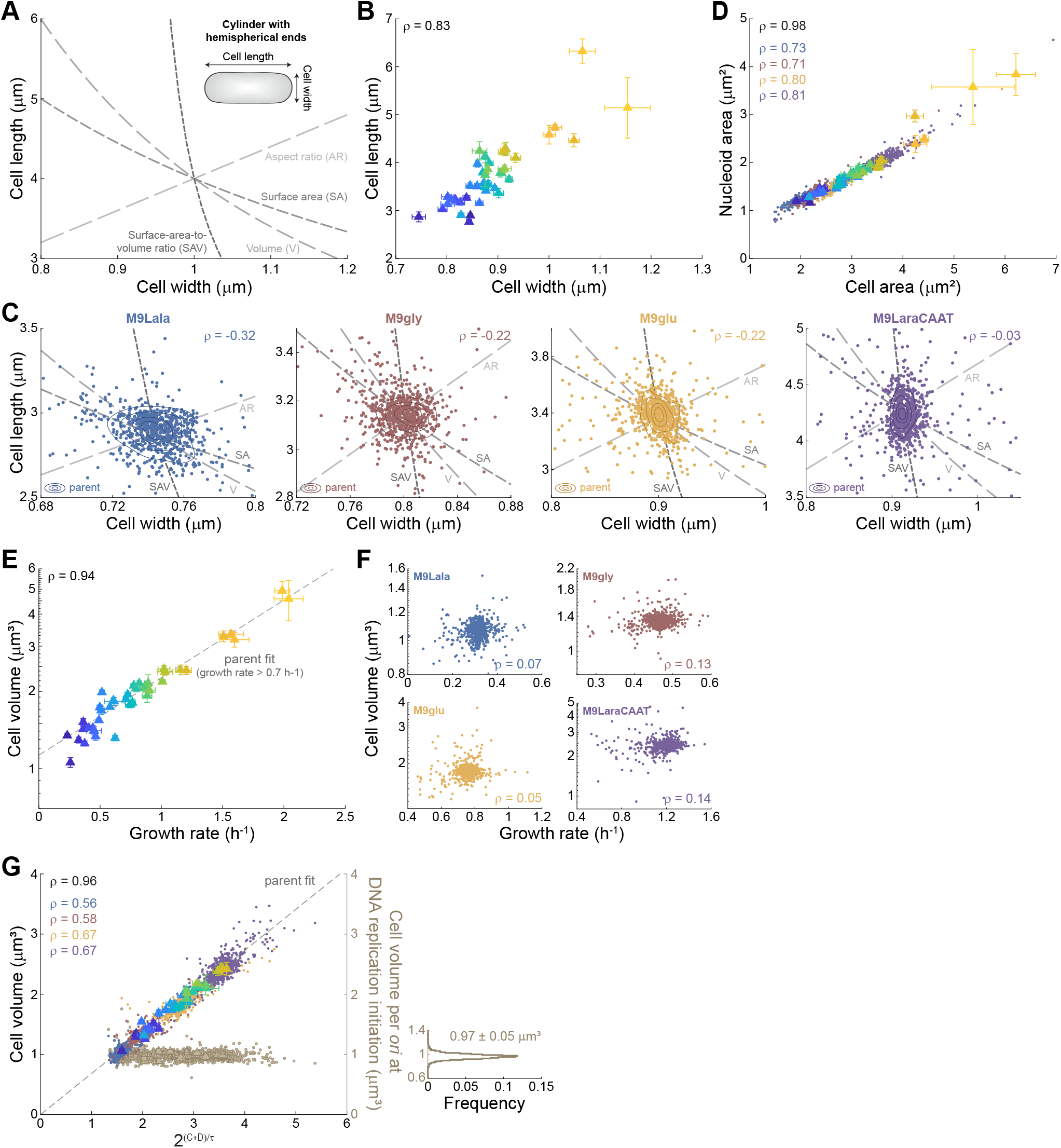
Examination of previously reported relationships. In all panels, triangles show the mean values of the indicated features for the parent strain across nutrient conditions (color-coded based on Figure 1B), with error bars indicating SD between individual experiments (n ≥ 4). Small dots indicate the data for the deletion strains, colored according to the nutrient condition. The Spearman correlation coefficient (ρ) is shown for each colored data, with ρ in black showing the correlation for the parent strain data. A. Predicted relationships between cell length and width if cells (with an initial cell length of 4 μm and cell width of 1 μm) were to maintain a constant aspect ratio, surface area, volume, or surface-area-to-volume ratio. Cells were modelled as cylinders with hemispherical ends (inset). B. Scatter plot of mean cell width versus mean cell length for the parent strain across nutrient conditions. C. Scatter plots of mean cell width versus mean cell length for the gene deletion strains. Each panel shows a different nutrient condition. Isocontours represent the 0.10, 0.25, 0.50, and 0.75 probability envelopes of the parent replicates in the same nutrient condition. Shown in grey are the predicted relationships between cell length and width if cells were to maintain an aspect ratio (AR), surface area (SA), volume (V), or surface-area-to-volume (SAV) ratio equal to that of the parent strain in the same nutrient conditions. Panel axes were cropped to remove outliers and more clearly show most of the data. D. Scatter plot of mean nucleoid area versus mean cell area for both parent and deletion strains across nutrient conditions. E. Scatter plot of mean cell volume versus growth rate for the parent strain across nutrient conditions. The dotted grey line shows an unconstrained linear fit for growth rates > 0.7 h^-1^. F. Scatter plots of mean cell volume versus growth rate for the deletion strains in each nutrient condition. G. Scatter plot of mean cell volume versus the average number of *ori* (given by 2^(C+D)/τ^). The dotted grey line shows a linear fit of the parent data across nutrient conditions (with the intercept constrained to the origin). Scatter plot of the mean cell volume per *ori* at initiation of DNA replication versus the average number of *ori* is shown in beige. The frequency distribution of cell volumes per *ori* at the initiation of DNA replication, combining all measurements, is shown on the right. Mean ± SD is indicated on the plot. See also Figure S3.

Across nutrient conditions, it is well known that both cell length and width increase with increasing growth rate^5,46,48^, leading to a positive correlation between the two variables (Figure 2B). However, we did not observe this positive relationship across deletion strains, irrespective of the nutrient condition (Figure 2C). The Spearman correlation (ρ) between cell length and width was nonexistent in the nutrient-rich medium (M9LaraCAAT), consistent with our genome-wide study^34^, and weakly negative in the nutrient-poorer media (M9Lala, M9gly, and M9glu). Importantly, regardless of the growth medium, the data collectively failed to follow any specific relationship (shown by the dashed lines in Figure 2C) that would argue for cell size sensing through surface area, volume or the ratio of the two. These findings illustrate the importance of sampling large numbers and different types of perturbations to test general relationships.

### Nucleoid length robustly scales with cell length

Whereas the relationship between cell length and width widely differed across strains and nutrient conditions, other features exhibited more consistent behavior. In agreement with a scaling relation between nucleoid and cell size^34,49,50^, we observed a strong positive correlation between nucleoid and cell areas across all data (Figure 2D). Interestingly, nucleoid length displayed a stronger linear relationship with cell length than nucleoid width with cell width (Figure S3A-B), suggesting that nucleoid length scaling is the main driver behind the overall size scaling of the nucleoid.

### DNA replication initiates at a constant mean cell volume per *ori*

Our dataset also allowed us to revisit debated bacterial laws. First, we confirmed a recent report^21^ that the nutrient growth law (i.e., the exponential scaling between cell volume and growth rate) becomes less reliable in nutrient-poor conditions (α < 0.7 h^-1^) with cells becoming smaller than predicted based on their growth rates (Figure 2E). In addition, we observed no correlation between cell volume and growth rate for the deletion strains within the four nutrient conditions (Figure 2F), which is consistent with our previous study^34^. Growth rate was also uncorrelated with other morphological variables such as cell surface area and surface-area-to-volume ratio (Figure S3C-D).

On the other hand, the recently challenged general growth law in which cell volume scales linearly with the average number of *ori* (given by 2 ^(C+D)/τ^)^14,15^ did hold across all perturbations, with data points from both the parent and deletion strains collapsing onto the same curve (Figure 2G). In contrast, the proposed alternative between cell volume and α * (C + D)^21^ did not yield the expected linear relationship (Figure S3E). The agreement of our data with the general growth law indicates that the mean cell volume per *ori* at DNA replication initiation remains largely constant across strains and nutrient conditions (0.97 ± 0.05 μm^3^, mean ± SD, n = 3137 perturbations) (Figure 2G). Only the cell volume at which *E. coli* initiates DNA replication per *ori*, and not cell length, area, or surface area (Figure S3F), was generally constant across strains and conditions. This indicates that, at the population level, initiation of DNA replication in *E. coli* is coupled to cell volume and no other cell morphological feature, similar to in *Bacillus subtilis^51^*.

### The relative timing of cell cycle events negatively correlates with cell size

To examine other potential dependencies, we plotted the relative cell cycle timings of the parent strain versus its average cell volume or growth rate across nutrient conditions. For all cell cycle events but FtsZ ring formation (see below), we found a strong negative, linear dependency with cell volume (Figure 3A) and, to a lesser extent, with growth rate (Figure S3G). These linear relationships are valuable for their predictive power. They enable the inference of any cell cycle timing (initiation of cell constriction, initiation and termination of DNA replication, initiation of nucleoid constriction, and separation of nucleoids) for the parent strain based on a simple cell volume measurement of the population in any given nutrient condition. Similarly, determination of a single relative timing (e.g., initiation of cell constriction) allows inference of not only the average cell volume, but also the timing of any other cell cycle event (e.g., termination of DNA replication or nucleoid separation).

**Figure 3.**
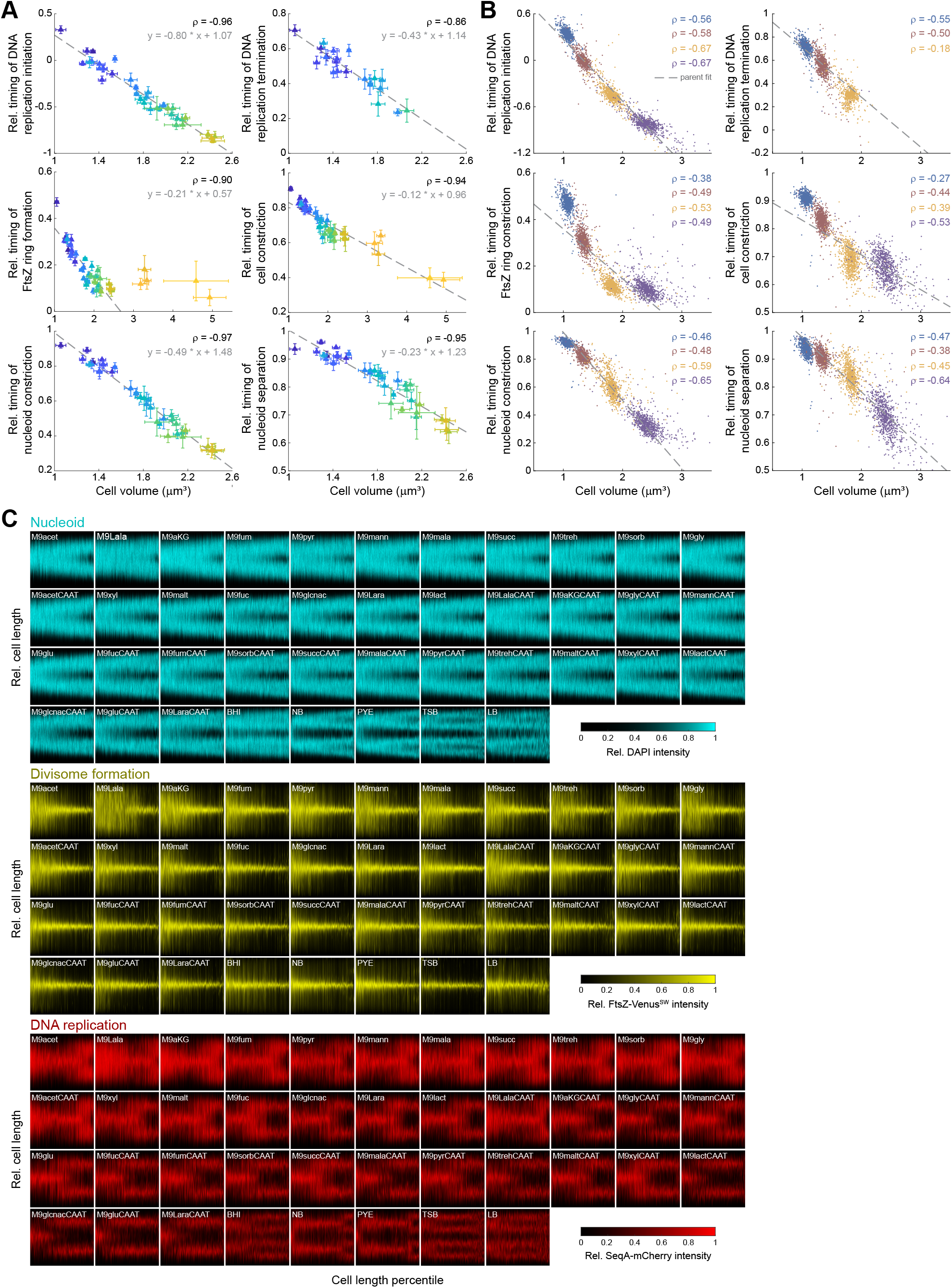
Cell size dependence of the timing of cell cycle events. A. Scatter plots of the relative timing of the indicated cell cycle events versus mean cell volume for the parent strain across nutrient conditions (color-coded based on Figure 1B), with the error bars indicating SD between individual experiments (n ≥ 4). ρ = Spearman correlation coefficient. The dotted grey line indicates the unconstrained linear fit of the parent strain data across nutrient conditions. For FtsZ ring formation, the linear fit was constrained to exclude the five richest nutrient conditions (LB, TSB, PYE, NB, and BHI). For these five conditions, timings for DNA replication and nucleoid constriction or separation could also not be reliably extracted (see Methods). The relative timing of DNA replication termination could not be determined in rich nutrient conditions in which cells displayed overlapping rounds of DNA replication (see Methods). B. Same as panel A but for the deletion strains across nutrient conditions. Colors indicate the nutrient condition in which strains were imaged. The dotted grey lines correspond to the linear fits for the parent data shown in panel A. Note that we were unable to extract timings of DNA replication termination for deletion strains in M9LaraCAAT (data usually in purple) due to the presence of overlapping DNA replication forks. C. Demographs of DAPI, FtsZ-Venus^SW^, and SeqA-mCherry signals in the parent strain across the indicated nutrient conditions, showing nucleoid dynamics, divisome formation and DNA replication characteristics, respectively. Cell length is normalized to show the relative cell length in the y-axis. Nutrient conditions were ordered based on increasing growth rate.

However, these empirical relationships appeared to only apply to metabolic changes, as the mutant data deviated to varying degrees (Figure 3B). Interestingly, deletion strains with larger cells tended to execute cell cycle events earlier in the cell cycle, as shown by the negative ρ values across deletion strains under the same nutrient condition (Figure 3B). Since such a negative correlation was not observed for growth rate (ρ ~ 0, Figure S3H), these results suggest a form of cell size control for all examined cell cycle events.

To examine this connection with cell size, we tested all pairwise correlations between each cell cycle event’s relative or absolute timing and all other features across strains and nutrient conditions. We also looked at whether the observed cell size correlation originates from a sizer-, adder- or timer-like behavior. For sizer-like behavior, we expected the relative or absolute timing of the cell cycle event to occur at a specific average cell or nucleoid size across strains and media. For cell cycle events under control of an adder, we expected them to occur after cells had added, on average, a constant length/area/surface area/volume since an earlier event. Finally, for timer-like behavior, the cell size control would be indirect through a dependence on an earlier cell cycle event that would itself be under cell size control. This could originate from a constant time delay (i.e., timer) between the two events, corresponding to the time needed for the cell to prepare the downstream event (e.g., divisome assembly if the downstream event were cell constriction). Such a systematic delay could either scale with the cell doubling time (i.e., relative timer) or be invariable across growth rates (i.e., absolute timer), depending on whether the underlying preparatory process is growth rate-dependent or not. In some cases, we could not reliably extract some cell cycle timings in the richest media (see Methods). Despite this limitation, our dataset remained rich in variability as evidenced by the wide range of absolute and relative timings obtained for each cell cycle event (Figure 3A-B). Furthermore, when needed, we used demographs of the DAPI, SeqA-mCherry, and FtsZ-Venus^SW^ signals to extract qualitative information (see below). Such demographs consist of linear representations of integrated fluorescence of cells sorted by their length^52^. They provide an overview of DNA replication, nucleoid and FtsZ dynamics along the cell division cycle across media (Figure 3C).

Our goal was to identify cell cycle laws, i.e., design principles that apply across strains and nutrient conditions. From this extensive analysis emerged four new potential laws, one per cell cycle event discussed below.

### Nucleoid constriction occurs at a constant mean nucleoid length

In our study, the onset of nucleoid constriction marked the first visible step of bulk chromosome segregation. This step varied in timing across our dataset, ranging from 0.2 to 0.9 in relative cell cycle time and ~10 min to > 2.5 h in absolute time (Figure 4A). Within this broad range, nucleoid constriction occurred at a relatively constant nucleoid length (3.20 ± 0.25 μm, n = 3260 perturbations, Figure 4B), consistent with a sizer principle for the initiation of nucleoid constriction. Nucleoid constriction also tended to initiate at a constant cell length (3.89 ± 0.31 μm, Figure 4C) given the scaling between nucleoid length and cell length (Figure S3A).

**Figure 4.**
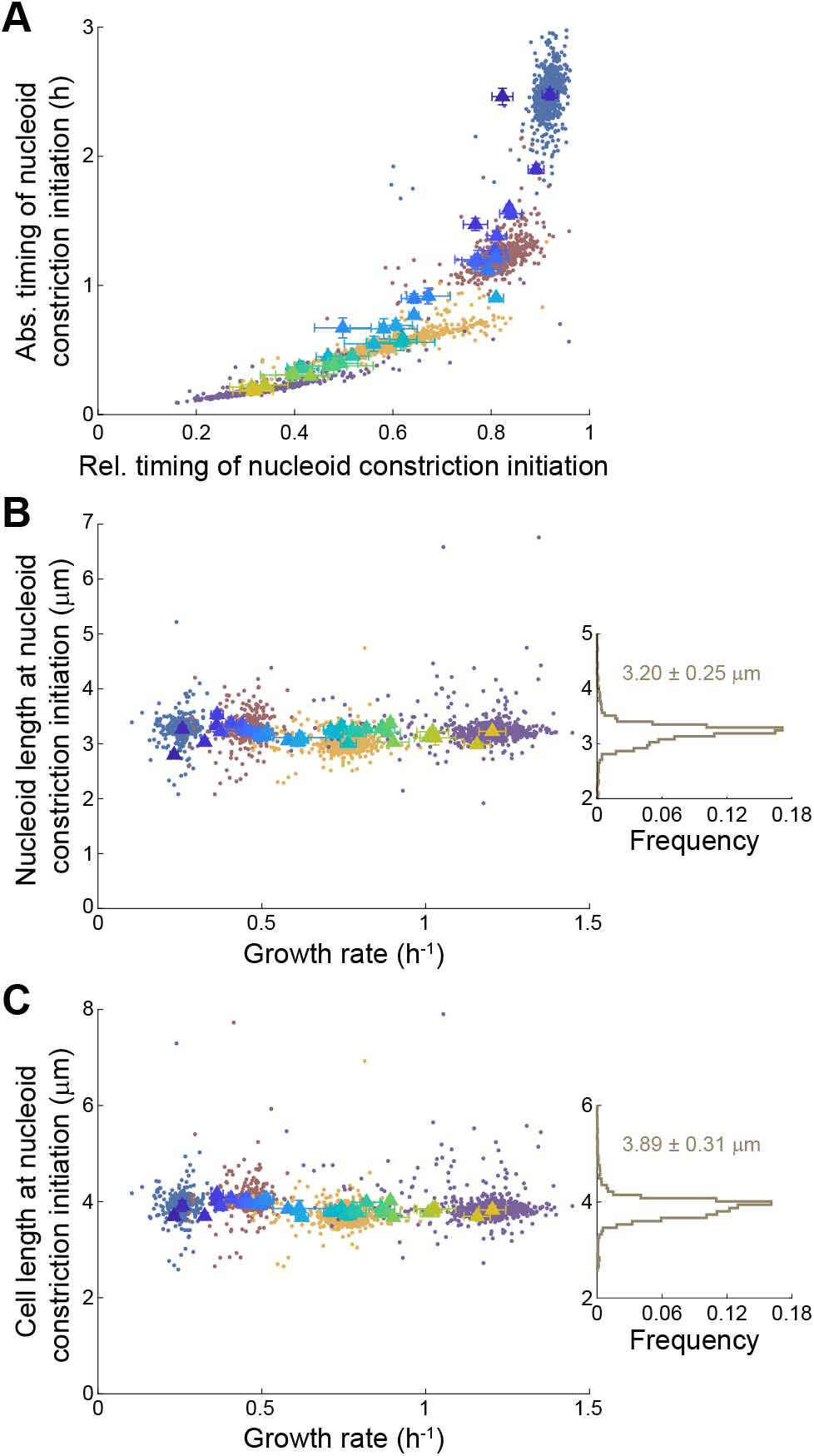
Constant nucleoid length at nucleoid constriction. In all panels, triangles show the mean values of the indicated features for the parent strain across nutrient conditions (color-coded based on Figure 1B), with the error bars indicating SD between individual experiments (n ≥ 4). Small dots indicate the data for the deletion strains, colored according to the nutrient condition. The Spearman correlation coefficient (ρ) is shown for each colored data, with ρ in black showing the correlation for the parent strain data. Initiation of nucleoid constriction could not be reliably determined in the five richest nutrient conditions (LB, TSB, PYE, NB, and BHI) because it occurred in the previous generation. A. Scatter plot of the absolute timing of nucleoid constriction initiation from birth versus its relative timing. B. Scatter plot of the nucleoid length at nucleoid constriction initiation versus growth rate. The frequency distribution of nucleoid lengths at nucleoid constriction initiation, combining all measurements, is shown on the right. Mean ± SD is indicated on the plot. C. Scatter plot of the cell length at nucleoid constriction initiation versus growth rate. The frequency distribution of cell lengths at nucleoid constriction initiation, combining all measurements, is shown on the right. Mean ± SD is indicated on the plot.

### The order of cell cycle events changes with the nutrient quality

Next, we examined cell division events, starting with the earliest one we observed: FtsZ ring formation. Unlike DNA-related events, FtsZ ring formation only showed a negative correlation with cell volume in nutrient-poor conditions (Figure 3A). This was because FtsZ ring formation is limited to one per division cycle, as illustrated in FtsZ-Venus^SW^ demographs of the parent strain across nutrient conditions (Figure 3C). These demographs confirmed that the accumulation of FtsZ-Venus^SW^ at midcell, which marks the formation of the FtsZ ring, occurs later in poor media relative to richer media. However, even in the richest nutrient conditions (Figure 3C), these demographs showed no clear evidence of additional FtsZ rings at the ¼ and ¾ positions (which would become midcell positions in the future daughter cells). This indicates that the number of FtsZ rings is constrained to one per generation. Because of this “generational constraint”, the relative timing of FtsZ ring assembly at midcell reached a plateau near the beginning of the cell cycle in media where populations have a mean cell volume ≳ 2 μm^3^ (Figure 5A). In contrast and as expected, initiation of DNA replication could often be traced back to the previous generation in richer nutrient conditions (Figure 5A). Analysis of the parent strain demographs also revealed that, in the five richest conditions, DNA replication even initiated two generations prior (as evidenced by the appearance of four separate signals; Figure 3C). Similarly, nucleoid separation could also occur in a previous generation. This was showcased in DAPI demographs of the parent strain grown in the richest nutrient conditions, where cells were already born with two separated nucleoids (Figure 3C), as described before^53^.

**Figure 5.**
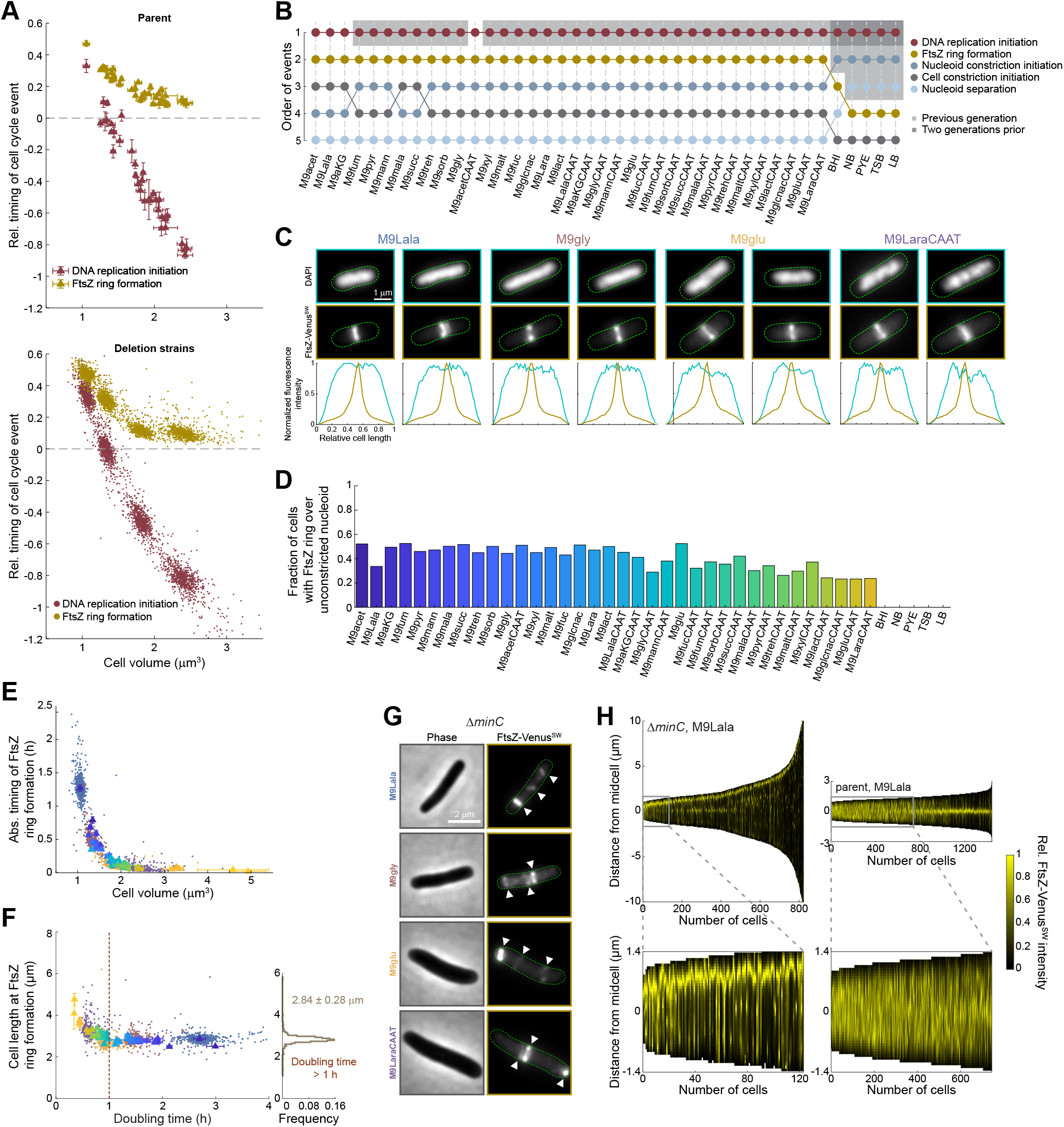
Generational constraint and cell length threshold of FtsZ ring formation. A. Scatter plot of the relative timing of FtsZ ring formation and DNA replication initiation for the parent strain (top) and deletion strains (bottom) versus mean cell volume across nutrient conditions. Error bars indicate SD between individual experiments (n ≥ 4). Negative values indicate that the cell cycle event occurred one or two generations prior. B. Relative order of cell cycle events across nutrient conditions for the parent strain. Nutrient conditions were ordered based on increasing growth rate. Shadings indicate cell cycle events that occurred one (light grey) or two generations (dark grey) prior. C. Top, representative fluorescence images of the parent strain in the indicated nutrient conditions. Bottom, fluorescence intensity profiles of DAPI and FtsZ-Venus^SW^ signals for these cells. D. Bar graph showing the fraction of cells with a FtsZ ring over an unconstricted nucleoid across nutrient conditions (ordered based on increasing growth rate). E. Scatter plot of the absolute timing of midcell FtsZ ring formation versus cell volume. Triangles show the mean values of the parent strain across nutrient conditions (color-coded based on Figure 1B). Error bars indicate SD between individual experiments (n ≥ 4). Small dots indicate the data for the deletion strains, colored according to the nutrient condition. F. Scatter plot of cell length at FtsZ ring formation versus doubling time. Triangles show the mean values of the parent strain across nutrient conditions (color-coded based on Figure 1B). Error bars indicate SD between individual experiments (n ≥ 4). Small dots indicate the data for the deletion strains, colored according to the nutrient condition. The frequency distribution of cell lengths at FtsZ ring formation, for strains and nutrient conditions giving rise to a doubling time > 1 h, is shown on the right. Mean ± SD are indicated on the plot. G. Representative phase contrast and FtsZ-Venus^SW^ images of Δ*minC* cells in the indicated nutrient conditions. White arrows indicate probable FtsZ rings. H. Top, demographs of FtsZ-Venus^SW^ signals of the Δ*minC* and parent strains in M9Lala. Bottom, zoomed-in parts of the demographs providing a more detailed picture of FtsZ-Venus^SW^ behavior in cells whose length falls below the identified cell length threshold of ~2.8 μm. See also Figure S4.

The absence of a generational constraint for DNA-related events resulted in a change in the order of cell cycle events when populations reached a mean cell volume of ~3 μm3 (growth rate of ~1.5 h^-1^). FtsZ ring formation at midcell switched from preceding nucleoid constriction and separation to following these DNA-related events in the richest media where these events happened in the previous generation (Figure 5B). Similarly, because of its dependence on FtsZ ring formation, cell constriction changed from almost coinciding with nucleoid constriction (Figure S4A) and from preceding nucleoid separation to becoming the last cell cycle event to occur in the richest media (Figure 5B).

### A single FtsZ ring forms after cells reach a certain length

The switch in event order was surprising given the paradigm that a FtsZ ring cannot form until after the nucleoid has constricted to create DNA-free space at the middle of the cell^54–56^, a phenomenon often referred to as nucleoid occlusion. Outside the five richest media, FtsZ ring formation preceded nucleoid constriction in the other 36 tested media (Figure 5B), leading to the emergence of many cells (> 30%) with a FtsZ ring over an unconstricted nucleoid (Figures 5C-D). Thus, nucleoid occlusion does not appear to be a robust inhibitory mechanism for FtsZ ring formation across nutrient conditions.

While FtsZ ring formation occurred almost immediately following cell birth in rich media associated with mean cell volumes > 2 μm3 (doubling time > 1 h), the timing of this cell cycle event varied dramatically in nutrient-poorer conditions, from almost right away to over 2 h after cell birth (Figure 5E). This variability was not due to differences in FtsZ amount or concentration within the cells, as these variables displayed little to no correlation with the relative or absolute timing of FtsZ ring formation (Figure S4B). Instead, in these nutrient-poorer conditions (doubling time > 1 h), we found that FtsZ ring formation occurred at a relatively constant cell length (2.84 ± 0.28 μm, n = 1583 perturbations; Figure 5F). Together with the observation of the generational constraint (Figures 3C, 5A and 5E), these results suggest that FtsZ ring formation follows a two-rule principle: 1) cells must have reached a certain length threshold and 2) there can only be one FtsZ ring per division cycle.

The Min system alone could be responsible for implementing both rules (see Discussion). This system, which includes the FtsZ polymerization inhibitor MinC^57^, has been extensively studied for its role in FtsZ ring positioning along the cell length^58,59^. In the absence of MinC, bacterial cells form minicells and filaments due to the division septum forming near the cell poles^60,61^. Consistent with the role of the Min system in constraining FtsZ to a single ring per cell, we observed cells with multiple FtsZ rings in the *E. coli ΔminC* mutant strain throughout all sampled nutrient conditions (Figure 5G). In addition, this strain already formed (polar) FtsZ rings in small cells, at sizes significantly smaller than the length threshold below which no FtsZ rings were observed in the parent strain (Figure 5H).

### Cell constriction occurs after a constant mean cell length has been added

FtsZ ring formation is often explicitly or implicitly considered as the main control point for cell division and, by extension, cell size determination. This raises the question of whether FtsZ ring formation triggers a temporal sequence of biochemical events (divisome assembly) that leads to cell constriction. Therefore, we examined whether there was a constant absolute or relative timer between FtsZ ring formation and cell constriction. Relative or absolute timers between two events can be identified by plotting the relative or absolute timing of these events across strains and nutrient conditions. If a timer existed between two cell cycle events, they would display a positive linear relationship with a slope of 1 and an intercept (x = 0) corresponding to the time delay between the two events. We found no evidence of a timer between FtsZ ring formation and the onset of cell constriction (slope ≠ 1; Figure S5A). Instead, the relative and absolute time delays between these events varied across conditions (Figure 6A-B). Similar observations were made for the relationship between FtsZ ring formation and the completion of cell division (Figure 6CD).

**Figure 6.**
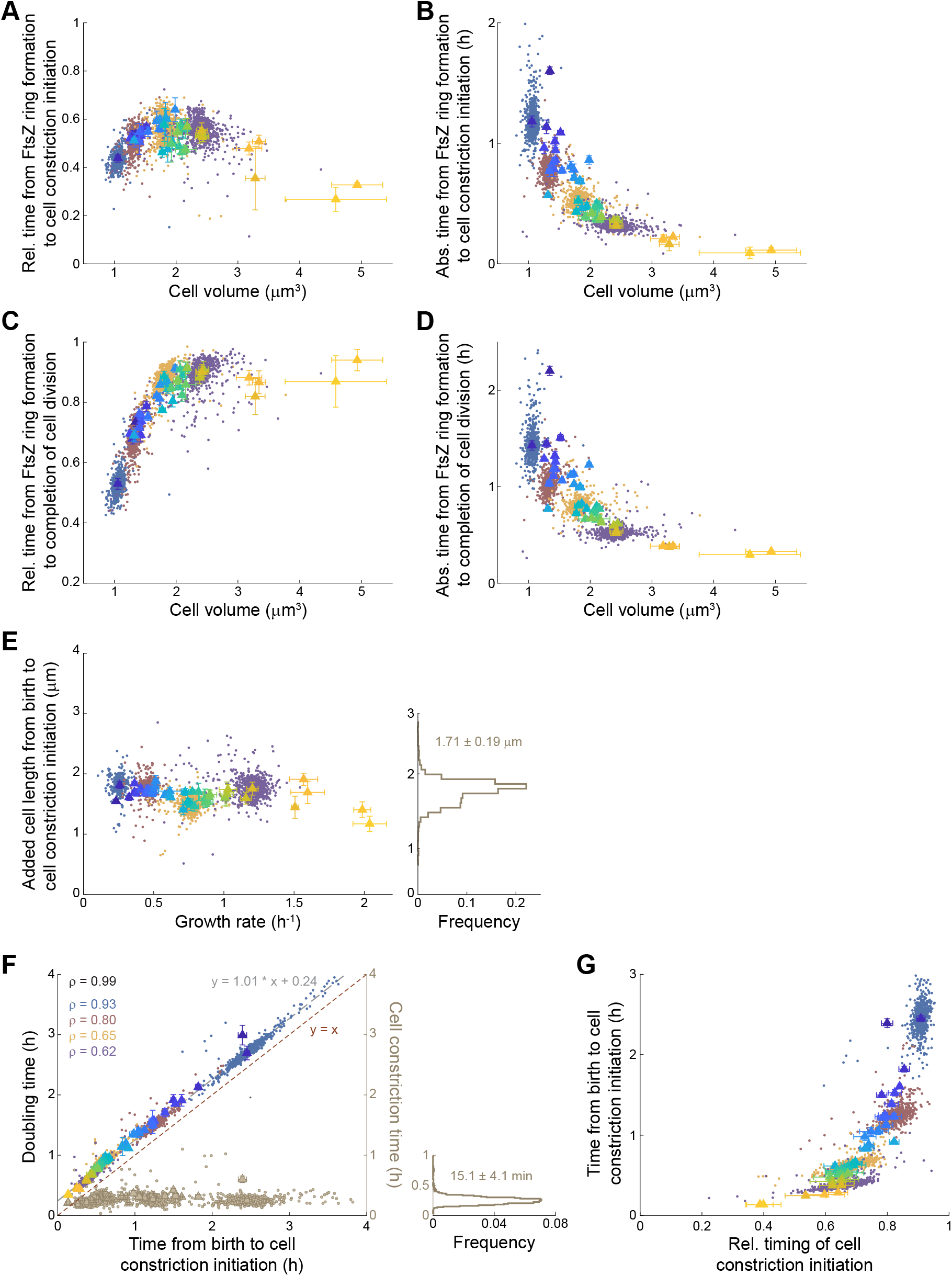
Cell constriction as a control point for cell size determination. In all panels, triangles show the means of the indicated features for the parent strain across nutrient conditions (color-coded based on Figure 1B), with the error bars indicating SD between individual experiments (n ≥ 4). Small dots indicate the data for the deletion strains, colored according to the nutrient condition. A. Scatter plot of the relative time from FtsZ ring formation to cell constriction initiation versus cell volume. B. Scatter plot of the absolute time from FtsZ ring formation to cell constriction initiation versus cell volume. C. Scatter plot of the relative time from FtsZ ring formation to completion of cell division versus cell volume. D. Scatter plot of the absolute time from FtsZ ring formation to completion of cell division versus cell volume. E. Scatter plot of the added cell length from birth to cell constriction initiation versus growth rate. The frequency distribution of added cell lengths from birth to cell constriction initiation, combining all measurements, is shown on the right. Mean ± SD are indicated on the plot. F. Scatter plot of doubling time versus the time from birth to cell constriction initiation. The dotted grey line indicates the unconstrained linear fit of all data across strains and nutrient conditions. The dotted red line represents the bisector (y = x), where data points would collapse if both events coincided. The Spearman correlation coefficient (ρ) is shown for each colored data, with ρ in black showing the correlation for the parent strain data. Scatter plot of the time from initiation of cell constriction to completion of cell division (i.e., the cell constriction time) versus the time from birth to cell constriction initiation is shown in beige. The frequency distribution of cell constriction times, combining all measurements, is shown on the right. Mean ± SD are indicated on the plot. G. Scatter plot of the absolute time from birth to cell constriction initiation versus the relative timing of cell constriction initiation. See also Figure S5.

For the initiation of cell constriction, the closest principle of constancy we identified across strains and nutrient condition was a relatively constant added cell length between birth and cell constriction (Figure 6E). The distribution of added lengths between birth and cell constriction was relatively narrow (1.71 ± 0.19 μm, n = 3166 perturbations) across strains and conditions compared to the wide range of average cell lengths (from ~3 to ~6 μm) (Figure 6E). This suggests an adder principle for the control of cell constriction initiation whereby, at the population level, cell constriction tends to occur after a constant mean cell length has been added after birth.

### Cell division follows cell constriction after a constant time delay

The closest timer-like behavior (i.e., least variable time delay) we observed between two cell cycle events was an absolute timer between constriction initiation and cell separation. This was evident by i) the high positive correlation between cell constriction initiation and doubling time, and ii) the collapse of the data on a linear relationship with a slope of 1 and an intercept of 0.24 h (Figure 6F). This means that completion of cell division follows the initiation of cell constriction after an average time of ~15 min (n = 3083 perturbations; Figure 6F). There was some variability in this time delay, with a few outliers across nutrient conditions (e.g., parent strain in M9acet) and deletion strains (Figure 6F). Regardless, the distribution of the time window remained small (SD = 4.1 min) across strains and media compared to the wide range of doubling times (from 20 min to 4 h). As a result of this apparent timer, the timing of initiation of cell constriction happened early in the cell division cycle of fast-growing strains and late in slow-growing ones (Figure 6G). This timer-like behavior also argues that the initiation of cell constriction is a key control point for cell size determination (see Discussion).

### The nutritional environment predominantly contributes to the phenotypic landscape

While most deletion strains abide by the cell cycle laws proposed above, we identified mutants that significantly deviate from each law. There were two types of outliers depending on whether the mutant phenotype depended on the nutrient condition or not. The most common type was the nutrient-dependent outliers, which were defined by having *s* > 3 (or *s* < −3) for at least one nutrient condition and *s* < 1.5 (or *s* > −1.5) in at least one other condition (Figure 7A). In other words, these mutants broke the rule (i.e., deviated by at least three SD from the parent’s median value) under some nutrient conditions but either behaved similarly to the parent (typically) or deviated from the parent in the opposite direction (rarely) under other nutrient conditions. Note that, in this analysis, we did not consider the cell length threshold principle for FtsZ ring formation because phenotypic robustness could only be assessed under two, instead of four, nutrient conditions due to the generational constraint in the two richest media (Figure 5E-F).

**Figure 7.**
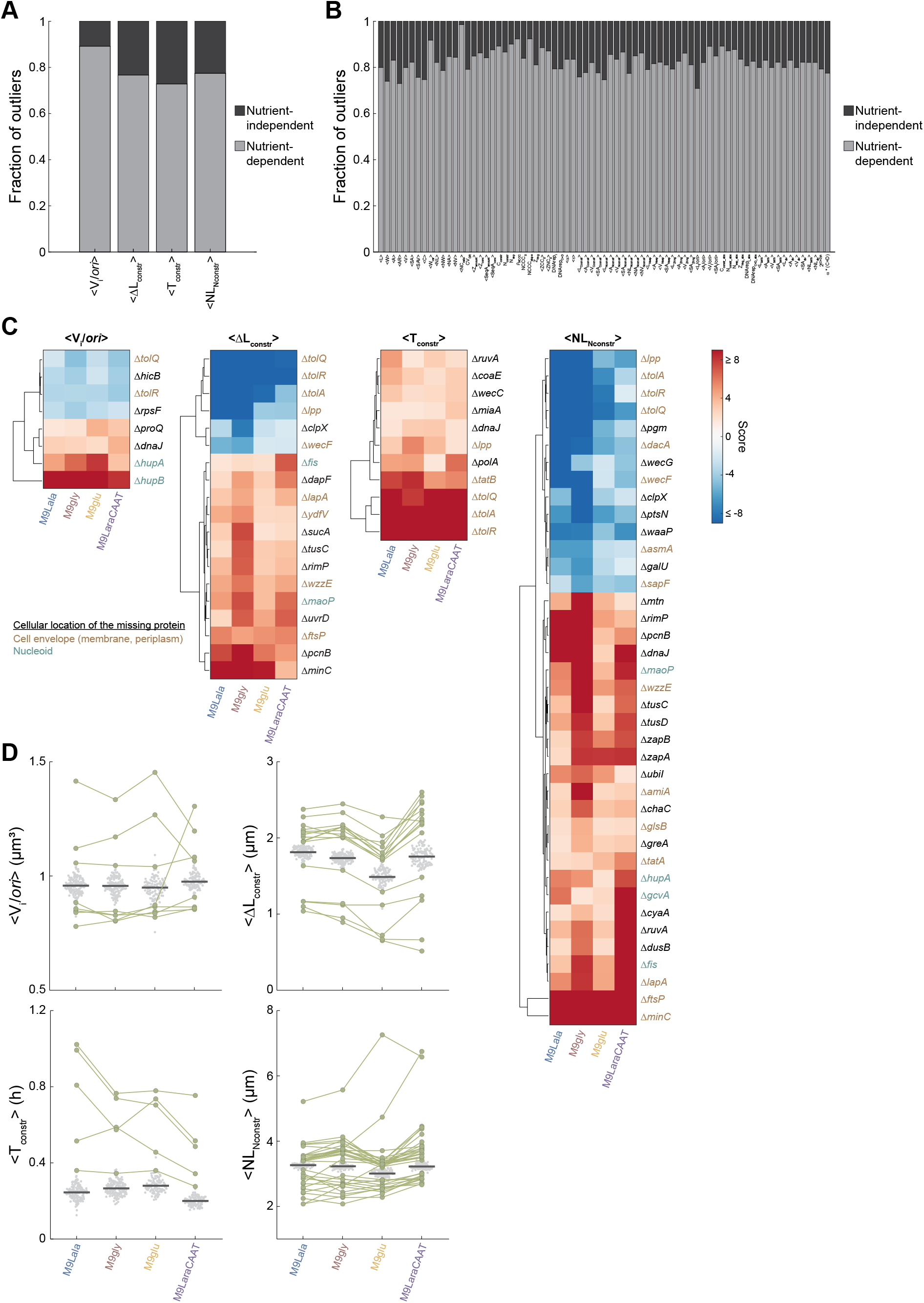
Rule breakers and nutrient-dependent phenotypic landscapes. A. Bar graphs showing the fraction of nutrient-independent and -dependent outliers for the following cell cycle laws: constant DNA replication initiation volume per *ori* (V_i_/*ori*), constant added cell length between birth and cell constriction (ΔL_constr_), constant time between initiation and completion of cell constriction (T_constr_), constant nucleoid length at initiation of nucleoid constriction (NL_Nconstr_). Nutrient-dependent outliers were defined by having *s* > 3 (or *s* < −3) for at least one nutrient condition but *s* < 1.5 (or *s* > −1.5) in at least one other condition. Nutrient-independent outliers were defined as having *s* > 3 (or *s* < −3) for at least one nutrient condition and *s* > 1.5 (or *s* < −1.5) in all other conditions. B. Bar graphs showing the fraction of nutrient-independent and -dependent outliers for 73 extracted population-level features (see Table S5 for the definition of symbols). Four features were omitted (DNArep_C_, DNArep_D_, DNArep_C_abs_, and DNArep_D_abs_) as these could not be reliably extracted in all four nutrient conditions (see Methods). C. Clustergrams showing the normalized scores of nutrient-independent outliers for the indicated cell cycle laws. D. Outliers (green) with |*s*| ≥ 2 across the four tested nutrient conditions for the indicated cell cycle laws. Parent replicates are shown as beeswarm plots in grey. Dark grey lines indicate the mean of parent replicates in each nutrient condition. See also Figure S6.

Nutrient-dependent phenotypes were not limited to the biological laws we identified. They were also common for all 77 features quantified in our study (Figure 7B). For example, for cell length, we identified 195 strains (24% of all gene deletion strains) that were significantly longer or shorter than the parent (*s* > 3 or *s* < −3) in at least 1 nutrient condition (expected false discovery rate of 1.2%), but the majority of them (156 strains) also displayed either parent-like cell lengths or deviations in the other direction (*s* < 1.5 or *s* > −1.5) in at least one other condition. A similarly large fraction of nutrient-dependent deviating phenotypes was observed for all other features (Figure 7B). To illustrate, Figure S6 shows an example of a nutrient-dependent phenotype with the Δ*clpX* strain (Figure S6A), which displayed minicell formation in nutrient-poor conditions (M9Lala and M9gly) but not in richer nutrient conditions (M9glu and M9LaraCAAT). These minicells, which are small anucleate cells produced by aberrant cell divisions at the cell poles^62^, were likely formed by perturbing the normal interaction between the ClpXP protease complex, the Min system and FtsZ^63,64^. In line with these observations, the Δ*clpP* strain displayed a similar and even more severe nutrient-dependent phenotype, from normally dividing in M9LaraCAAT to producing minicells in M9gly and M9glu to not growing in M9Lala (Figure S6A).

The predominance of nutrient-dependent phenotypes across features (Figure 7A-B) suggests an important role for the nutrient condition and, more broadly, central metabolism in modulating the integration of cell cycle progression with cell morphogenesis and growth rate.

### True rule breakers are rare

A small but important subset of outliers was nutrient-independent (Figures 7C). Many of the genes deleted in these strains encode proteins that localize to either the cell envelope or the nucleoid (Figure 7C), stressing the importance of these two structures in cell replication. Interestingly, among mutants that displayed a deviating phenotype across nutrient conditions (regardless of the direction), many still followed a rule of constancy reasonably well. Consider, for example, all mutants with an ļ*s*ļ > 2 for nucleoid constriction across the four tested media. Most of these mutants still initiated cell constriction at a similar mean nucleoid length (<NL_Nconstr_>) in all four nutrient conditions, only at a lower or higher value relative to the parent (Figure 7D). Only a few gene deletion strains broke the rule of constancy by displaying large differences in values across conditions (Figure 7D). The scarcity of true rule breakers demonstrates how robust the rules of constancy are to the cellular replication of *E. coli*.

## Discussion

### *E. coli* follows simple rules of cell cycle control at the population level

By phenotyping growth, cell cycle and cell morphological features across large-scale perturbations, we show that *E. coli* follows unifying principles of cell cycle control at the population level. We confirmed that, on average, *E. coli* initiates chromosome replication at a constant cell volume per *ori* (Figure 2G). In addition to this cell cycle law, our study suggests four others related to nucleoid segregation (Figure 4B) and different steps of cell division, including FtsZ ring formation (Figures 5E-F), initiation of cell constriction (Figure 6E), and completion of cell division (Figure 6F). We propose that *E. coli* uses these simple rules to integrate cell cycle progression with cell size and ensure robust cellular replication across many conditions.

### Cell cycle timings are connected to different aspects of cell size

The proposed cell cycle laws are all connected to some aspect of cellular morphology, explaining the observed cell size dependence of cell cycle events (Figure 3A-B). While cell size dependencies held across genetic perturbations in a given nutrient condition (Figure 3B), we observed near zero correlations between relative cell cycle timings and growth rate (Figure S3H). This argues that the cell cycle is coupled to cell size rather than growth rate. Growth rate is, in turn, integrated with cell size and the cell cycle via the general growth law (Figure 2G)^14,15^.

Interestingly, our results suggest that cell size dependence is achieved through different means depending on the cell cycle event. For instance, initiation of DNA replication is coupled to a constant cell volume per *ori* (Figure 2G), while FtsZ ring formation is connected to cell length (Figure 5F). Nucleoid constriction also appears connected to cell length (Figure 4B-C) by virtue of the scaling between nucleoid length and cell length (Figure S3A). For the initiation of cell constriction, we found no evidence for a cell size threshold (i.e., sizer) or a constant delay (i.e., timer) relative to other cell cycle events. Instead, our analysis is more consistent with a control that relies on an added cell length from cell birth (i.e., an adder) (Figure 6E). Completion of cell division then follows the onset of cell constriction after a timer (Figure 6F). Thus, in the case of the final step of cell division, the cell size-regulation occurs indirectly via a dependence on cell constriction.

Our observations are unlikely to be a peculiarity of *E. coli. B. subtilis* has also been shown to initiate DNA replication at a constant cell volume per *ori* at the population level^51^. Furthermore, this Gram-positive bacterium has been reported to display a constant ratio of FtsZ rings to cell length across four different growth rates^65^. This is consistent with a cell length threshold for FtsZ ring formation, suggesting that unifying rules in cell cycle control may be conserved.

### Population vs. single-cell principles

It is important to note that the proposed cell cycle laws consist of population-level principles obtained by averaging the behavior of many individual cells. As such, they reflect what *E. coli* has evolved to do on average as a system. At the single-cell level, these design principles could be achieved in different ways. For example, the constant average cell volume per *ori* at DNA replication initiation, which is observed at the population level, can theoretically be obtained either by a sizer at the single-cell level, where individual cells initiate DNA replication at a constant volume per *ori*, or by an adder, where individual cells add, on average, a constant volume per *ori* between consecutive DNA replication initiation events^66^. Time-lapse experiments have verified the adder scenario^44,67^.

Ultimately, combining knowledge from population and single-cell studies will be critical when searching for the biophysical mechanisms that underlie the cell cycle laws. In some cases, like the initiation of DNA replication at a constant cell volume per *ori*, there are theoretical and experimental clues pointing to potential cell size-dependent mechanisms^44,68–71^. In the case of FtsZ ring formation, the presence of a FtsZ ring over the nucleoid in most (36/41) tested media (Figures 5B-D) precludes a predominant role for nucleoid occlusion in controlling FtsZ ring assembly. On the other hand, the Min system alone may be sufficient to establish the two-rule principle regulating FtsZ ring formation: one ring per division cycle and the cell length threshold (Figure 5G-H). By tracking individual cells during growth, we showed in previous work that the oscillation amplitude of MinD (the partner protein of the FtsZ polymerization inhibitor MinC) depends on cell length^72^. In fact, the cell length threshold at which we observed FtsZ ring formation in this current study (~2.8 μm) falls within the length interval (2-3 μm) in which depletion of MinD/C (and thus the minimum of FtsZ inhibition at midcell) first became apparent in our previous work^72^. As for the generational constraint of FtsZ ring formation, it may emerge from the fact that the oscillation of MinD/C only produces multiple minima in long (≳9 μm) cells ^73^’^74^. Indeed, the generational constraint is abolished in the Δ*minC* mutant, as multiple FtsZ rings were able to form in single cells in all four tested media (Figure 5G-H).

We did not find evidence of robust timers across strains and media between cell cycle events, with the notable exception of the timer between cell constriction initiation and cell separation (Figure 6F). Early indications of this apparent timer already existed in the literature, although they were not discussed in this context. For example, a strong linear relationship between the doubling time and the timing of cell constriction from birth has been reported for *E. coli* cells across four nutrient conditions and three growth temperatures^75^. This relationship had a slope close to 1 (0.96) and an intercept of ~14 min (which is remarkably close to our value of ~15 min). A simple explanation for this timer would be that it reflects the time it takes for the biochemical process of cell constriction to occur. However, given that cells are wider in richer media, a constant time for cell constriction implies that wider cells constrict faster, as shown in Figure S5B. Consistent with this relationship, the cell constriction time has been shown to remain constant across a panel of MreB point mutants with varying cell widths^76^.

Importantly, the proposed cell cycle laws provide both benchmarks and constraints for testing current and future mechanistic models of cell cycle control. Furthermore, for each cell cycle law, our study identified gene deletion outliers (Figure 7), with the nutrient-independent ones being of particular interest. Some outliers do not break the rule per se; they only shift the constancy to a lower or higher set value (Figure 7D). While this underscores the importance of a constant value across conditions, it also renders the proteins missing in these mutant strains unlikely to be involved in mechanisms underpinning the laws. These outliers remain, however, valuable tools for testing hypotheses because they provide a high level of constraint, as any proposed mechanism should display the expected shift in control set points in these mutant strains. Our study also identified outliers for which the rule of constancy no longer applied (Figure 7D). In these mutants, the mechanism involved in a cell cycle law is likely broken, opening the door for future mechanistic studies.

### Points of control and integration

It is important to note that we extensively analyzed our dataset to examine all possible types of relationships (e.g., sizer, adder, timer) between the 77 features we measured. Only one design principle emerged for five cell cycle events, highlighting the discriminating aspect of our approach. In the literature, the bacterial cell cycle is often discussed in the context of two regimes: discrete vs. overlapping DNA replication cycles (slow vs. fast growth). Our findings suggest that this distinction may not be critical, as the proposed cell cycle laws apply regardless of whether DNA replication cycles are discrete or overlapping.

Throughout our analysis, we did not find any direct connection between DNA replication and division events across strains and media, supporting the idea that the DNA replication and cell division cycles operate largely independently. This is important as a wealth of models have been proposed to explain if and how cell division is controlled by DNA replication^13,21,39,44,53,66,67,77–80^. DNA replication and cell constriction are, however, indirectly connected by virtue of their dependence on (different aspects of) cell size, which may help explain the appearance of conflicting results across previous studies.

Our analysis allowed us to identify control points for cell cycle events. For example, while FtsZ and its polymerization into an apparent ring-like structure are required for the initiation (and therefore completion) of cell constriction, we did not find evidence of a constant timer, sizer, or adder that links cell constriction or cell division to a FtsZ feature (such as the timing of FtsZ ring formation or the concentration or abundance of FtsZ within the cell). Instead, we found support for a timer between the initiation of cell constriction and its completion. This constancy of cell constriction time suggests that the onset of cell constriction acts as a control point for cell size determination, given that the downstream events are executed within a fixed period. Similarly, we identified a constant nucleoid length as a likely control point for ensuring proper coordination of cell size with nucleoid segregation.

Interestingly, some nutrient-independent outliers for the proposed cell cycle laws broke more than one law (Figure 7). Most strikingly, we identified several *tol* mutants as consistent outliers for all five design principles (Figure 7C). TolQ, TolR and TolA are inner membrane components of the Tol-Pal system, which spans the entire cell envelope^81^. The Tol-Pal system is known to be involved in outer membrane invagination, septal peptidoglycan processing, and maintenance of outer membrane integrity^82–86^. A recent study identified *tol* mutants that initiated DNA replication at smaller cell volumes than the wild type^87^, which we confirmed (Figure 7C). Our study shows that gene deletions of individual Tol components can also lead to severe phenotypes related to the timing of FtsZ ring formation, cell constriction, nucleoid segregation, and completion of cell division (Figure 7C). The pleiotropic phenotypes of these deletion mutants (Figure S6B) suggest that this system could play a major role in integrating the DNA and division cycles.

### The carbon source often drives mutant cell morphology and cell cycle phenotypes

Whereas the motivation of our study was to test previously proposed relations and uncover design principles of cell cycle regulation, our large dataset also allowed us to explore the phenotypic landscape of *E. coli* in more depth. We previously found that a large fraction (~20%) of the non-essential *E. coli* genome directly or indirectly affects cell morphology and/or cell cycle characteristics in a single growth condition^34^. Here, by varying the carbon sources, we showed that phenotypes usually depend on the nature of the nutrient available in the environment, as a severe (|s| ≥ 3) phenotype in one condition was often not persistent across other nutrient conditions (Figures 7A-B). This indicates that central metabolism, which is dictated by nutrient availability, is the primary driver of phenotypes. Consistent with this idea, deletion mutants of central carbon metabolism exhibit diverse cell morphological alterations^88^. There are also practical implications of a metabolic dependence. Cell size or cell cycle mutants are often characterized under a single nutrient condition with the implicit assumption that the phenotypes would persist in most other conditions. Our work suggests that mutants may need to be examined through the lens of the metabolism that the cells experience.

### Evolutionary implications

The large variability in phenotypes covered in our screen (Figure S2A) illustrates the broad range of the phenotypic space available to *E. coli*. Consider, for example, cell volume or the relative timing of DNA replication initiation. Across the tested nutrient conditions, over 200 deletion strains (i.e., almost a quarter of the examined deletion strains) differed by three or more standard deviations from the parent. Yet, by and large, mutants follow all five proposed cell cycle laws (Figures 2G, 4B, 5E-F, 6E, and 6F), including those related to cell size or DNA replication. Thus, the phenotypic variability in cell size, shape, and cell cycle timings contrasts with the robustness of the cell cycle laws across genetic and metabolic perturbations. This suggests that the cell cycle laws are under strong evolutionary constraints while cell morphology can more rapidly evolve, explaining the rapid emergence of strains of the same species that vary substantially in cell shape and size^89^. The cell cycle laws provide a simple way to ensure faithful cell cycle progression by connecting cell cycle events to an aspect of cell size, even if cell morphology changes through mutations.

### Reducing biological complexity to simple design principles

Our approach highlights the importance of using a large number of diverse perturbations to confidently identify design principles. In our case, this enabled the uncoupling of cellular features that might otherwise appear to be dependent on each other, such as cell length and width (Figure 2B-C) or growth rate and cell volume (Figure 2E-F). Using large-scale genetic perturbations achieves this effect by offering discriminating power, as it averages out the specific effect associated with each perturbation and increases confidence in any identified dependencies^34–37^.

The cell cycle laws, together with the general growth law, provide simple and robust means of coordinating cell cycle progression with cell size and growth rate across various conditions. Thus, there appears to be an inherent robustness in cell cycle regulation, where even complex phenotypes can be deconstructed into a combination of simple but shared principles.

Our approach of simultaneously quantifying multiple features across a large collection of mutants under several nutrient conditions should be broadly applicable to the study of other aspects of bacterial physiology. Simple general principles may exist for other cellular processes such as gene expression, cell wall synthesis, metabolism, energy production/homeostasis or second messenger production. Many of these processes can be tracked intracellularly through different fluorescent labeling approaches or metabolic biosensors^90–92^. These tools, together with transposon or CRISPR interference libraries, open the door for similar large-scale, microscopy-based studies in a wide variety of species. Such an approach may reveal universal principles of cellular replication and physiology.

## Supporting information

Key resources table

## Acknowledgments

We thank Dr. Harold Erickson for sharing the FtsZ sandwich fusion used in this work. We would also like to thank the Jacobs-Wagner laboratory for fruitful discussions and critical reading of the manuscript. We also thank the members of the Govers laboratory for feedback on the manuscript. S.K.G. was partly funded by postdoctoral fellowships from the Belgian American Educational Foundation, United States (B.A.E.F.), the de Duve Institute, Belgium, and the F.R.S.-FNRS, Belgium. C.J.-W. is an investigator of the Howard Hughes Medical Institute.

## Author contributions

Conceptualization, S.K.G., M.C. and C.J.-W.; Methodology, S.K.G., M.C. and C.J.-W. Software, S.K.G. and M.C.; Formal Analysis, S.K.G.; Investigation, S.K.G. and B.T.; Data Curation, S.K.G..; Writing – Original Draft, S.K.G. and C.J.-W.; Writing – Review & Editing, S.K.G., M.C., G.L., and C.J.-W; Visualization, S.K.G. and C.J.-W.; Supervision, C.J.-W.; Project Administration, C.J.-W.; Funding Acquisition, C.J.-W.

## STAR Methods

### RESOURCE AVAILABILITY

#### Lead contact

Further information and requests for resources and reagents should be directed to and will be fulfilled by the lead contact, Christine Jacobs-Wagner.

#### Materials availability

All bacterial strains generated in this study are available from the lead contact.

#### Data and code availability

Microscopy data reported in this paper will be shared by Dr. Sander Govers upon reasonable request. All original code developed as part of this study has been deposited in the publicly accessible Github code repository (https://github.com/JacobsWagnerLab/published).

### EXPERIMENTAL MODEL AND SUBJECT DETAILS

#### Bacterial strains

Strain CJW6324 (MG1655 *seqA::seqA-mcherry ftsZ::ftsZ-venus^SW^*)^49^ was used as the parent strain throughout this study. Individual deletions (n = 806) were moved into the parent strain by P1 phage transduction to assemble a deletion strain library. To do this, lysates were prepared from the corresponding deletion strains of the *E. coli* Keio collection^93^. Strains for which the transduction failed were constructed using lambda red recombineering^94^. After initial construction of the library, 115 deletion strains were randomly chosen and verified by colony PCR (primers are listed in Table S2). The small fraction (< 5%) of incorrect deletion strains identified by colony PCR were reconstructed using lambda red recombineering. While the FtsZ-Venus^SW^ fusion is known to cause a mild cell elongation phenotype particularly in nutrient-rich conditions^25,95^, it can support cell division as the sole source of FtsZ within a cell, and does so across all deletion strains included in this study. Any potential artefact of the fluorescent fusions used in this study is also mitigated by the fact that all strains are equipped with them, thereby enabling a relative comparison.

Individual gene deletion strains (806) were selected using a previous genome-wide, image-based quantitative screen of the Keio *E. coli* single-gene deletion collection grown in M9gluCAAT^34^. Among the 806 strains imaged in our study, 93 of them were selected based on their phenotypes related to the positive correlation between nucleoid size and cell size (nucleocytoplasmic ratio), and the negative correlation between nucleoid size and the relative timing of nucleoid separation. These strains were identified as outliers for these relationships by calculating their distance to a linear fit obtained by including all strains. These 93 strains were combined with 713 strains that clustered together in specific morphology islands (2, 3, 6, 7, 8, 9, 11, 12, 13, 14, 16, 17, 18, 19, 20, 21, 22) or cell cycle islands (2, 3, 4, 7, 8, 9, 11, 12) identified in our initial screen^34^, to obtain the final collection of 806 deletion strains. All islands contained strains with similar phenotypes and only the selected islands contained strains that displayed deviations in more features than growth-related features alone (growth rate and OD_max_). Genes were associated with clusters of orthologous groups (COGs) as in our previous work^34^.

### METHOD DETAILS

#### Growth conditions

An overview of the 41 different nutrient conditions used in this study is provided in Table S1. Each strain was grown to stationary phase in the appropriate nutrient condition in pre-culture tubes (2 ml of growth media) in a shaking water bath at 37°C. A dilution (>1/10000) was then used to inoculate fresh growth medium (2 ml). The resulting cultures were allowed to grow until they reached an optical density at 600 nm (OD_600_) between 0.05 and 0.3 depending on the nutrient condition (Figure S2C) before sampling for microscopy. The absence of strong biases in average feature behavior across this OD_600_ window (Figure S2C) suggests that most strains were sampled during steady-state exponential growth.

#### Microscopy

Cells were imaged on 1% agarose pads supplemented with either M9 buffer (for M9-based nutrient conditions) or the appropriate growth medium (for the richest, undefined nutrient conditions). To stain the chromosomal DNA that forms the nucleoid, cells were incubated with 1 μg/ml 4’,6-diamidino-2-phenylindole (DAPI) for 15 min in their growth medium prior to sampling for microscopy. To avoid having to correct for temporal biases introduced by cells spending longer times on agarose pads^34^, at most two deletion strains or parent replicates were spotted (0.5 μl) on the same pad. Separated spots were then traced back under the microscope and imaging was performed within 5-10 min after spotting the cells on the pad.

Phase contrast and epifluorescence imaging was performed on a Nikon Eclipse Ti microscope equipped with a 1.45 NA phase contrast CFI Plan Apo DM Lambda 100x oil objective (Nikon), an Orca-Flash4.0 V2 142 CMOS camera (Hamamatsu), and a Spectra X light engine (Lumencor). The following Chroma filter sets were used to acquire fluorescence images: DAPI (excitation ET350/50x, dichroic T400lp, emission ET460/50 m), YFP (excitation ET500/20x, dichroic T515lp, emission ET535/30 m) and mCherry/TexasRed (excitation ET560/40x, dichroic T585lpxr, emission ET630/75 m). The microscope was controlled by the NIS-Elements AR software.

#### Growth curves

Growth curves for the parent and deletion strains in various nutrient conditions were obtained in 96-well plates using a Synergy2 microplate reader (BioTek). Stationary phase precultures were inoculated into 150 μl fresh medium at a dilution of 1/300. Cultures were then grown for 60 h at 37°C during which OD_600_ measurements were taken every 4 min.

### QUANTIFICATION AND STATISTICAL ANALYSIS

#### Image processing

Individual cells were detected using the open-source software package Oufti^96^. Cell contours were curated in an automated fashion using a trained support vector machine model^49^ to automatically remove poor and incorrect cell detections. Cellular dimensions of retained cell contours were then quantified. Nucleoid outlines were detected using the objectDetection module in Oufti, from which nucleoid characteristics were extracted. The same parameters were used for all cell and nucleoid detections (see Tables S3 and S4 for the Oufti parameter values used in our study) to allow reliable comparison between strains and nutrient conditions. This is important, as the estimation of cellular features can vary with the chosen parameters^49^.

Note that a small consistent downshift of the added length value between birth and cell constriction initiation was observed in one nutrient condition (M9glu) relative to the other three conditions for both the parent and deletion strains (Figure 6E). This suggests the presence of a small systematic measurement error, though its origin is unknown.

The FtsZ-Venus^SW^ fusion provided information on the formation and position of the FtsZ ring^25,97^, whereas the SeqA-mCherry fusion allowed us to determine the DNA replication status of individual cells as well as the number of ongoing rounds of DNA replication^13,26,98^. FtsZ-Venus^SW^ and SeqA-mCherry signal information was added to Oufti-generated cell lists using the MATLAB function Add_FtsZ_Characteristics.m and Add_SeqA_Characteristics.m, respectively.

#### Extraction of cellular characteristics

Properties of individual cells (cell and nucleoid dimensions, DAPI fluorescence intensity, fluorescent marker behavior, etc.) were extracted from Oufti cell lists using the MATLAB function Extract_Extended_Cell_Properties.m. Morphological features (e.g., cell length, width, area, and volume) were determined by summing the dimensions of each segment of the cell mesh identified by Oufti (assuming cylindrical shape for the cell volume). See https://oufti.org/ for more details. These features of individual cells were used to calculate 77 different population features for each strain in each nutrient condition.

#### Extraction of average relative timings of cell cycle events

To extract the relative timings of cell cycle events from exponentially growing populations, we relied on an established method that uses the proportions of cells at different cell cycle stages to infer the relative cell ages at which specific cell cycle events occur under steady-state conditions^19,27,28^. This method has been experimentally validated through comparison with timelapse experiments by several groups^30–33^. In addition, for a previous study^34^, we performed a similar validation by comparing the relative timing of two cell cycle events (initiation of cell constriction and nucleoid separation), calculated using either time-lapse data or static images (i.e., randomly chosen frames from the same experiment). We found that the results from the time-lapse and static data were indeed in excellent agreement (see Figure on page 6 of the Peer Review Summary of our previous work^34^, which can be found at this weblink: https://www.embopress.org/action/downloadSupplement?doi=10.15252/msb.20177573&file=msb177573.reviewer_comments.pdf).

To calculate the relative timing of cell and nucleoid constriction, the fraction of unconstricted cells or nucleoids was determined using a constriction threshold of 0.2 μm (Figure S1B). This threshold was chosen conservatively to exclude only cells that had definitively committed to cell or nucleoid constriction. Importantly, constriction was measured in absolute length units to preclude cell width deviations from affecting the determination of constricting cells or nucleoids. For the relative timing of nucleoid separation, the fraction of cells with a single nucleoid was determined. For the relative timing of midcell FtsZ ring formation, the fraction of cells that did not display a clear FtsZ ring in the middle of the cell was determined using thresholds on the relative area and the relative midcell enrichment of the FtsZ-Venus^SW^ signal (Figure S1B). The midcell enrichment was assessed by comparing the FtsZ-Venus^SW^ signal in the middle 20% of a cell versus that in the remaining 80% of the cell.

Calculations of the relative timing of DNA replication initiation and termination relied on combining the SeqA-mCherry signal with cell volume measurements. In nutrient conditions in which cells displayed at least one clear period without ongoing DNA replication (as in M9Lala, M9gly, and M9glu media), as evidenced by a larger relative SeqA-mCherry area (Figure S1C, top row), distinct groups of cells could be identified (using k-means clustering), with each group corresponding to cells in a different DNA replication period. Depending on the DNA replication period, these cells differ in their number of origins (*ori*) and termini (*ter*) (Figure S1C). This information was used to calculate the average number of origins (<*ori*>) and termini (<*ter*>), which enabled the extraction of the relative duration of the C and D periods (Figure S1C)^14^. The relative duration of the B period, which corresponds to the relative timing of DNA replication initiation, is given by B = 1 - (C+D). In richer nutrient conditions (e.g., M9LaraCAAT), cells continuously replicated their DNA, as evidenced by the consistently small relative SeqA-mCherry area (Figure S1C, bottom row). In these conditions, termination of DNA replication could not be determined as other rounds of DNA replication were still ongoing. However, DNA replication re-initiation events were identified by examining the relative position of the (brightest) SeqA-mCherry focus, which moved from midcell to a quarter cell position upon re-initiation (Figures 3C and S1C). In contrast to how it has occasionally been interpreted^68,79^, this re-localization did not appear to be preceded by a termination event, as cells without ongoing DNA replication could not be detected (based on the relative area of the SeqA-mCherry signal; Figure S1C) in these nutrient-rich conditions. The presence of overlapping rounds of DNA replication, where a new round initiates before the previous one has terminated, is also in line with what is known about chromosome replication in *E. coli* in richer growth conditions (supporting growth rates > ~1 h^-4^)^8^. Combining the information on SeqA-mCherry localization with cell volume measurements again allowed the identification of distinct groups of cells (using k-means clustering) that corresponded to cells that differ in their number of *ori* (Figure S1C). We used this information to determine the average number of origins (<*ori*>) and extract the relative duration of the C+D period (Figure S1C).

The above-described methodology could not be used for the parent strain in the 5 richest conditions (LB, TSB, PYE, NB, and BHI), in which timings for DNA replication (because of the presence of more than two overlapping rounds of DNA replication) or nucleoid constriction (because it occurred in the previous generation) could not be reliably extracted.

#### Extraction of cell sizes at different cell cycle events

Assuming that individual cells grow exponentially at the single-cell level (in terms of cell or nucleoid length, area, volume, and surface area), the cell size at different cell cycle events was calculated based on the relative timing of a cell cycle event and the average cell size, as described previously^19^.

#### Reproducibility and effect of kanamycin

To verify the reproducibility of our experiments and assess the potential effect of the presence of kanamycin in the growth media, we imaged the original gene deletion mutants selected from the Keio collection (i.e., prior to phage transduction of the gene deletion into the parent strain expressing the FtsZ-Venus^SW^ and SeqA-mCherry markers) in M9gly with and without kanamycin (50 μg/ml). As before^34^, we found a strong correlation between cell length and width across duplicates, indicating good reproducibility (Figure S1D). For cell length, the presence of kanamycin led to a slight but consistent increase in cell length (Figure S1D). Therefore, for our study, we opted to grow and image the 806 deletions strains carrying both FtsZ-Venus^SW^ and SeqA-mCherry markers without kanamycin in the different growth media. Since the kanamycin resistance marker (which marks the gene deletion of each strain) is chromosomally encoded and is inserted in a gene for which the coding sequence is largely deleted^93^, we do not anticipate that the absence of kanamycin in the growth medium will have any detrimental effect to data interpretation.

#### Growth rate measurements

Growth rates were extracted from growth curves after well-specific blanking^99^. Blank values were obtained by averaging the first three optical density measurements of an experiment (during which cultures had not yet reached the detectable range of the microplate reader). The maximal growth rate was extracted from the corrected growth curves by fitting the Gompertz function^100^. To account for plate-to-plate variability, we set the median growth rate values for each plate in a given nutrient condition to the median growth rate value of the parental strain in that same nutrient condition.

#### Normalized score calculations

For each population-level feature in one of the four nutrient conditions in which we sampled our gene deletion mutant library, we also calculated normalized scores (*s*):

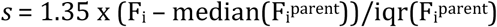

where F_i_ is the measured value of feature *i* in a nutrient condition, F_i_^parent^ are the values of the parent strain for that feature in the same nutrient condition and iqr is the interquartile range. We used this calculation based on the median and the interquartile range as it is more robust to outliers (compared to a typical z-score calculation). Given that the interquartile range of normally distributed data is equal to 1.35 times their standard deviation, we scaled the score by this factor to express the scores in terms of standard deviations away from the median of the parent values.

#### Demographs

Demographs, which are linear representations of integrated fluorescence of cells sorted by their length^52^, provide an overview of fluorescence signal dynamics from cell birth to division. Demographs in Figure 3C were constructed using the MATLAB function normLDemographOriented.m. Demographs in Figure 5H were constructed within Oufti.

#### Correlation coefficients

Spearman correlation coefficients (ρ) between variables were calculated using MATLAB’s built-in corr function.

#### Unconstrained linear fits

Unconstrained linear fits were performed using MATLAB’s built-in polyfit function.

#### K-means clustering

K-means clustering was performed sing MATLAB’s built-in kmeans function.

#### False discovery rate calculation

The expected false discovery rate of incorrectly identifying a deletion strain as one with a deviating phenotype (|*s*| > 3) for a single feature in at least 1 nutrient condition was calculated using the binomial formula 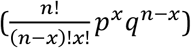.

## Supplemental figure legends

**Figure S1.**
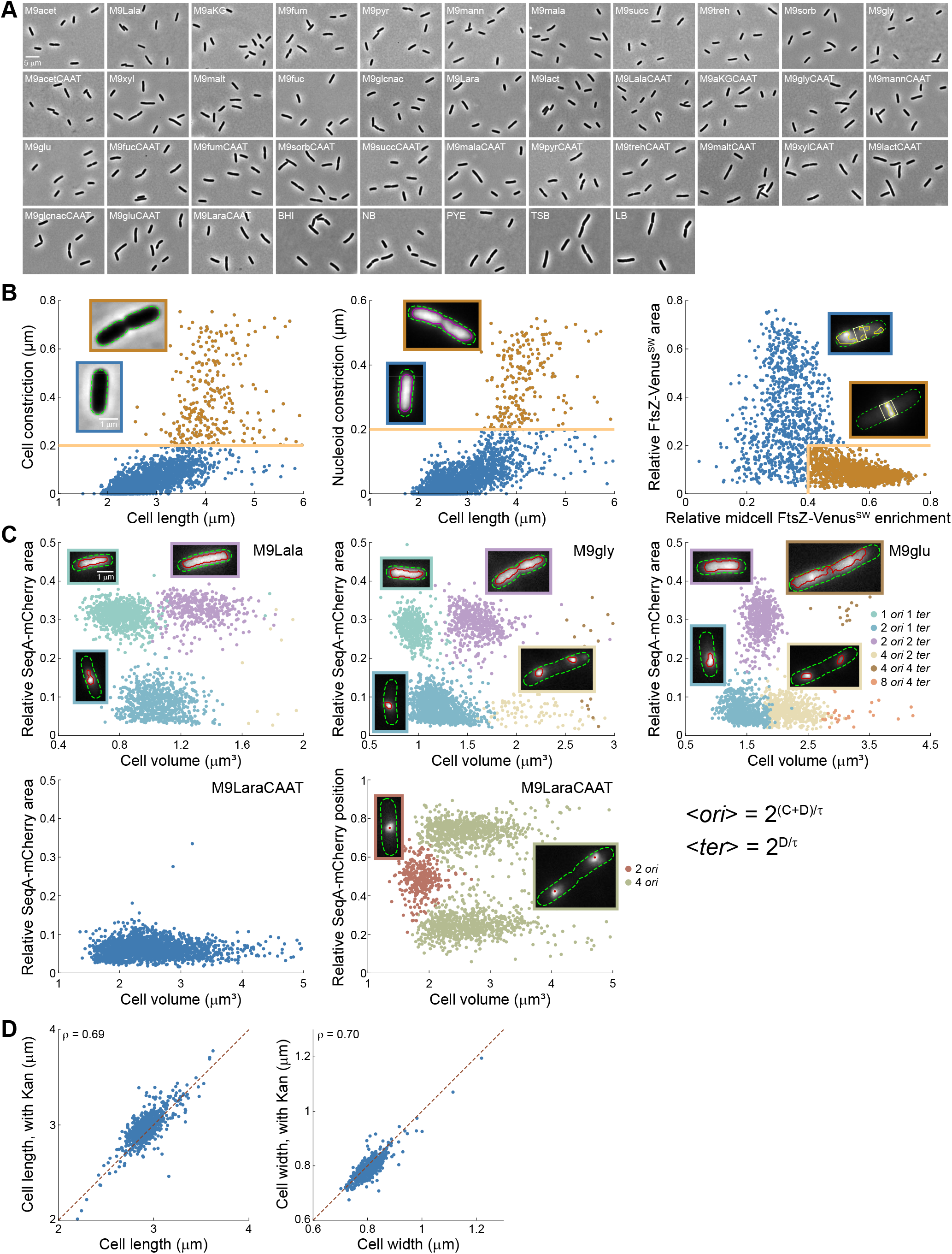
Extraction of cell cycle timings across nutrient conditions, related to Figure 1. A. Representative phase contrast images of the parent *E. coli* strain grown in liquid cultures of the indicated growth media at 37°C. See Table S1 for a full description of growth media. B. Determination of the fraction of unconstricted cells, unconstricted nucleoids, and cells without a FtsZ ring. Cell (green) and nucleoid (magenta) contours were generated using Oufti. FtsZ areas were detected using additional analysis scripts (see STAR Methods). C. Identification of cells in different DNA replication periods across nutrient conditions. In nutrient environments that did not give rise to overlapping DNA replication rounds, scatter plots of the relative SeqA-mCherry signal area versus cell volume were used to identify cells in different DNA replication periods (using k-means clustering). This information was then used to calculate the average number of origins and termini of replication, which enabled the extraction of the relative duration of the C and D periods. In richer nutrient conditions that gave rise to overlapping rounds of DNA replication, scatter plots of the relative position of the brightest SeqA-mCherry focus versus cell volume were used to identify cells in different DNA replication periods. This information was then used to calculate the average number of origins of replication, which enabled the extraction of the relative duration of the C+D period. D. Scatter plots showing the correlation in either mean cell length or mean cell width between replicates of the 806 original gene deletion mutants (lacking the FtsZ-Venus^SW^ and SeqA-mCherry markers) grown in M9gly with and without kanamycin. The dotted red line represents the bisector (y = x). The Spearman correlation coefficient (ρ) is shown for each plot.

**Figure S2.**
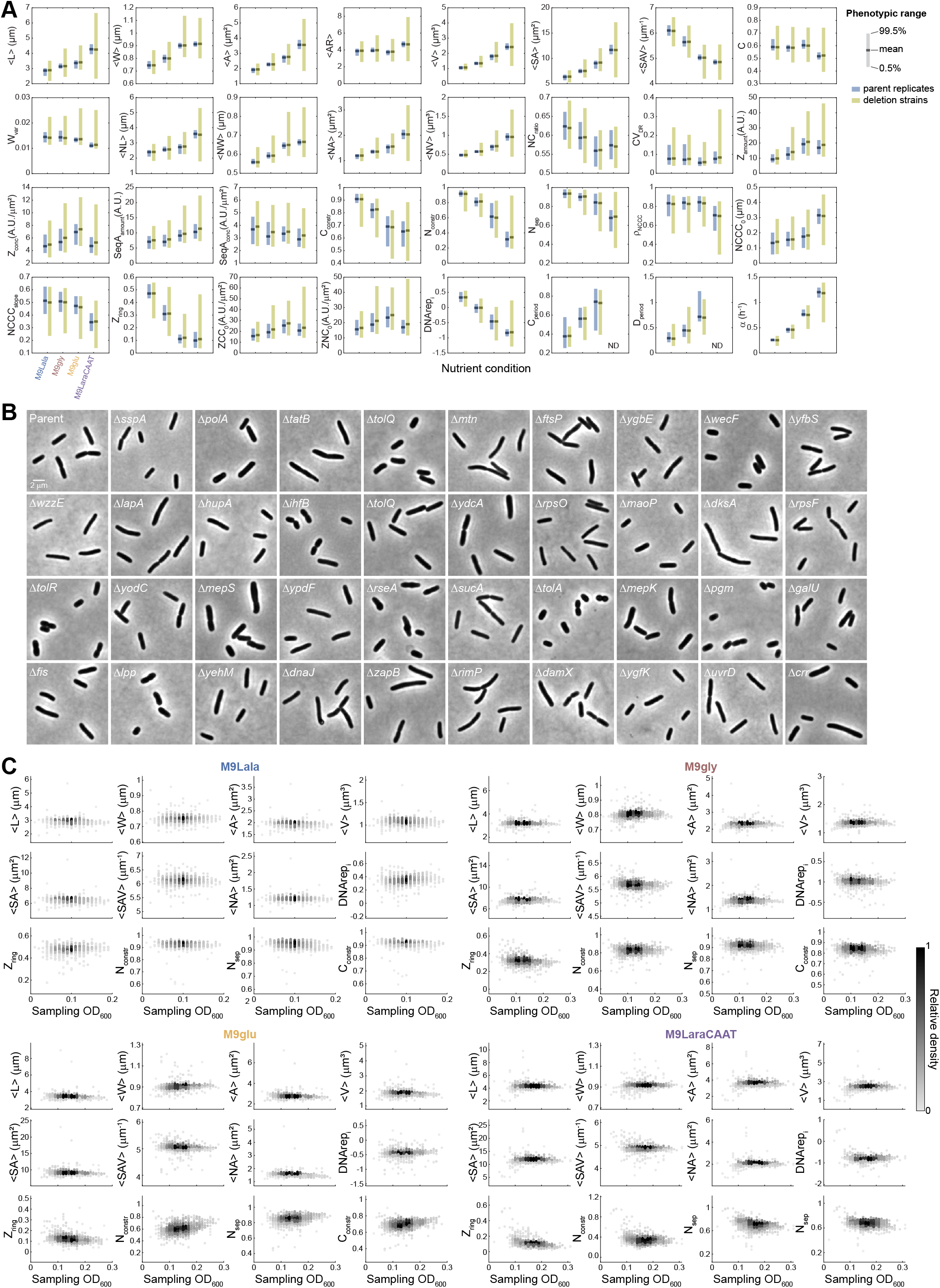
Phenotypic diversity and sampling effects, related to Figure 1. A. Bar plots indicating the phenotypic range (0.5-99.5% of the data) for extracted morphological, cell cycle, and growth rate features across nutrient conditions. Blue and green indicate the parent replicates and the deletion strains, respectively. Horizontal lines indicate the mean. ND indicates features that could not be determined. B. Representative phase-contrast images of the parent strain and indicated deletion strains grown in M9gly. C. Density scatter plots of morphological, cell cycle, and growth rate features across the optical density (OD_600_) window at which cells were imaged.

**Figure S3.**
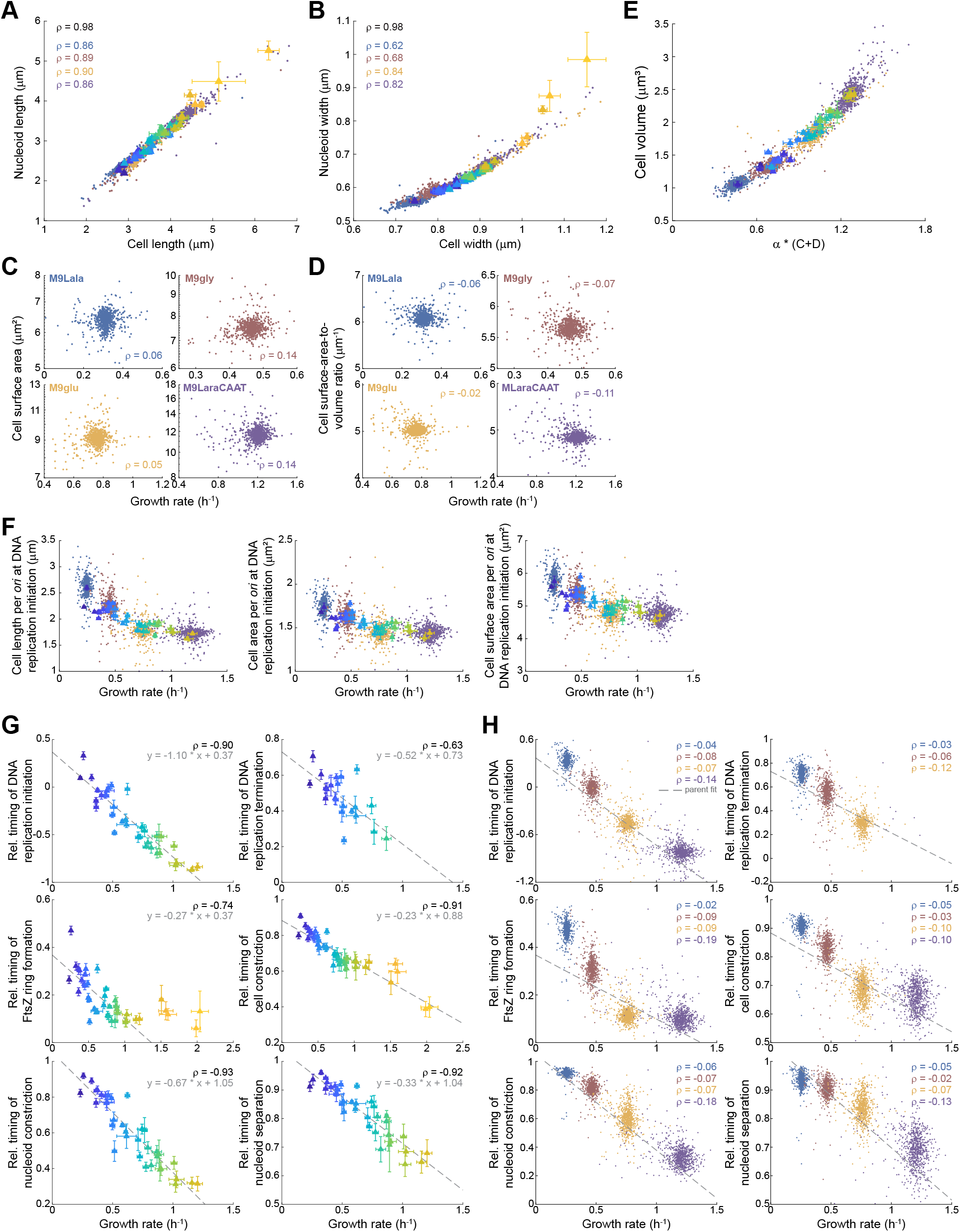
Relationships between growth rate and cell morphological or cell cycle features, related to Figures 2 and 3. In all panels, triangles show the means of the indicated features for the parent strain across nutrient conditions (color-coded based on Figure 1B), with the error bars indicating SD between individual experiments (n = ≥ 4). Small dots indicate the data for the deletion strains, colored according to the nutrient condition. When relevant, the Spearman correlation coefficient (ρ) is shown for each colored data, with ρ in black showing the correlation for the parent strain data. A. Scatter plot of mean nucleoid length versus mean cell length. B. Scatter plot of mean nucleoid width versus mean cell width. C. Scatter plots of cell surface area versus growth rate for the deletion strains. Each panel shows a different nutrient condition. D. Same as panel C, but for cell surface-area-to-volume ratio versus growth rate. E. Scatter plot of cell volume versus α*(C+D), with α = growth rate and C+D = the absolute duration of the C+D period. F. Scatter plots of the cell length, area, or surface area per *ori* at the initiation of DNA replication versus growth rate. G. Scatter plots of the relative timing of indicated cell cycle events versus growth rate for the parent strain across nutrient conditions. The dotted grey line shows an unconstrained linear fit. H. Same as panel F, but for the deletion strains across nutrient conditions. The dotted grey lines correspond to the linear fits for the parent data shown in panel G. Note that we were unable to extract timings of DNA replication termination for deletion strains in M9LaraCAAT (data usually in purple) due to the presence of overlapping DNA replication forks.

**Figure S4.**
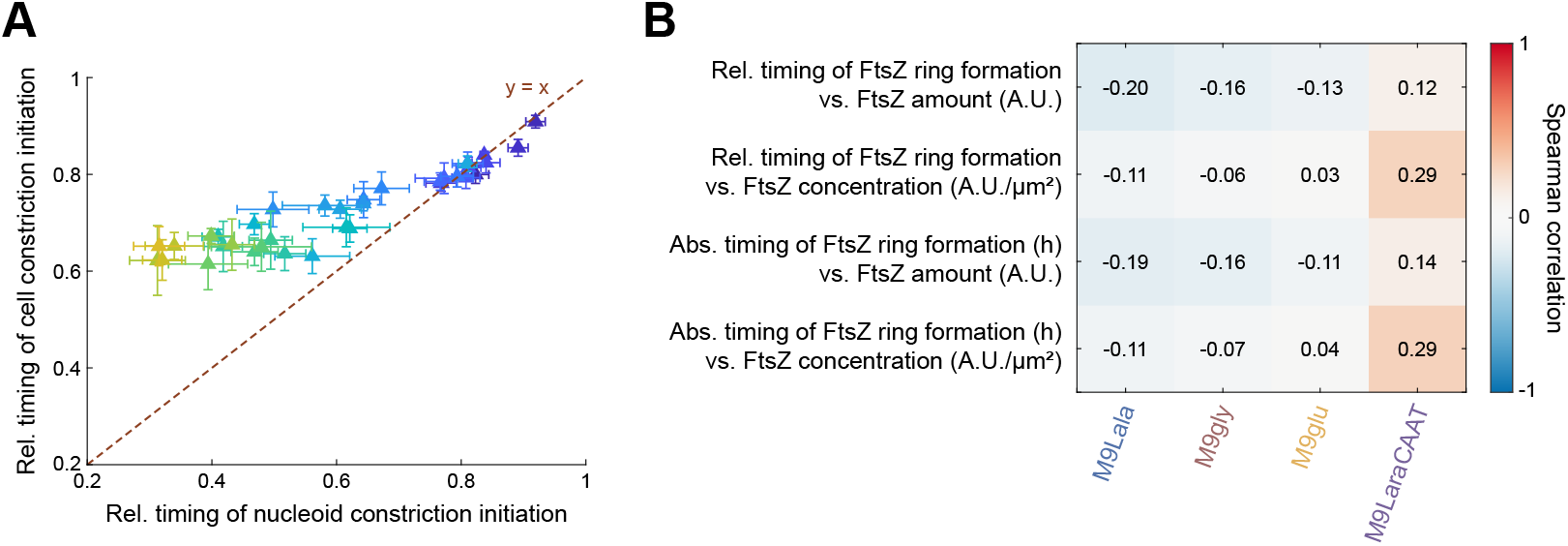
Relationships between nucleoid constriction, cell constriction and FtsZ levels, related to Figure 5. A. Scatter plot of the relative timing of cell constriction initiation versus the relative timing of nucleoid constriction initiation for the parent strain across nutrient conditions, with the error bars indicating SD between individual experiments (n = ≥ 4). The dotted line represents the bisector (y = x), where data points would collapse if both events coincided. This is the case in nutrient-poor conditions (darker blue colors), which explains the occasional switch in relative order of these events in Figure 5B. Data from the five richest nutrient conditions (LB, TSB, PYE, NB and BHI) could not be included in this graph because nucleoid constriction initiation occurred in the previous generation in these conditions (see demographs in Figure 3C), and its timing could not be reliably determined using the method employed for this figure. B. Heatmap showing the Spearman correlation coefficient between the FtsZ amount or concentration and the relative or absolute timing of FtsZ ring formation across the indicated nutrient conditions.

**Figure S5.**
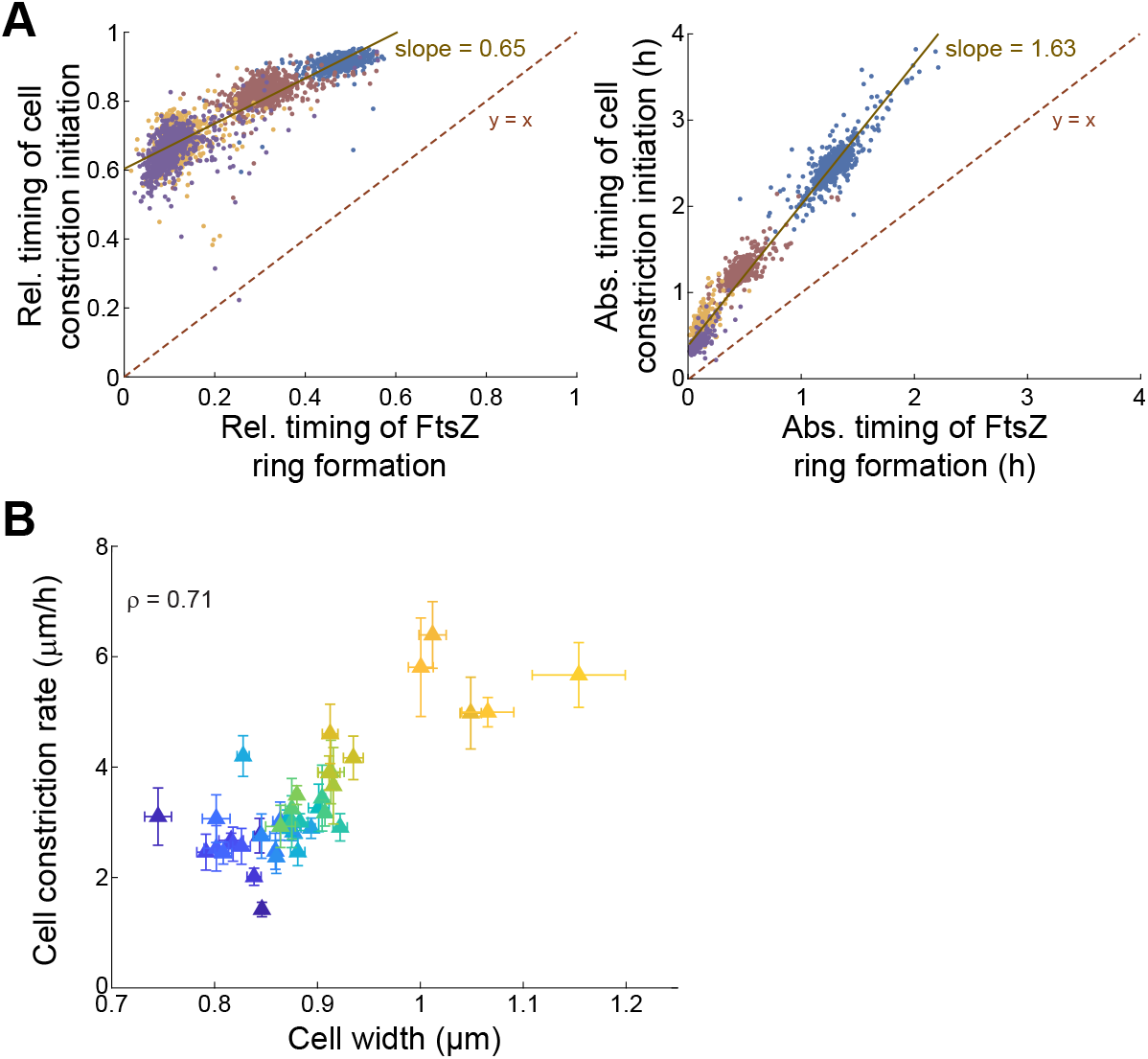
Relationships between FtsZ ring formation and cell constriction or between cell constriction rate and cell width, related to Figure 6. A. Scatter plots of the relative timing of cell constriction initiation since birth versus the relative timing of FtsZ ring formation (left) and the absolute timing of cell constriction initiation since birth versus the absolute timing of FtsZ ring formation (right) for the deletion strains across nutrient conditions. The dotted red line represents the bisector (y = x), where data points would collapse if both events coincided. Solid lines represent unconstrained linear fits across the data. B. Scatter plot of cell constriction rate versus mean cell width. Triangles show the means of the parent strain across nutrient conditions (color-coded based on Figure 1B), with the error bars indicating SD between individual experiments (n = ≥ 4).

**Figure S6.**
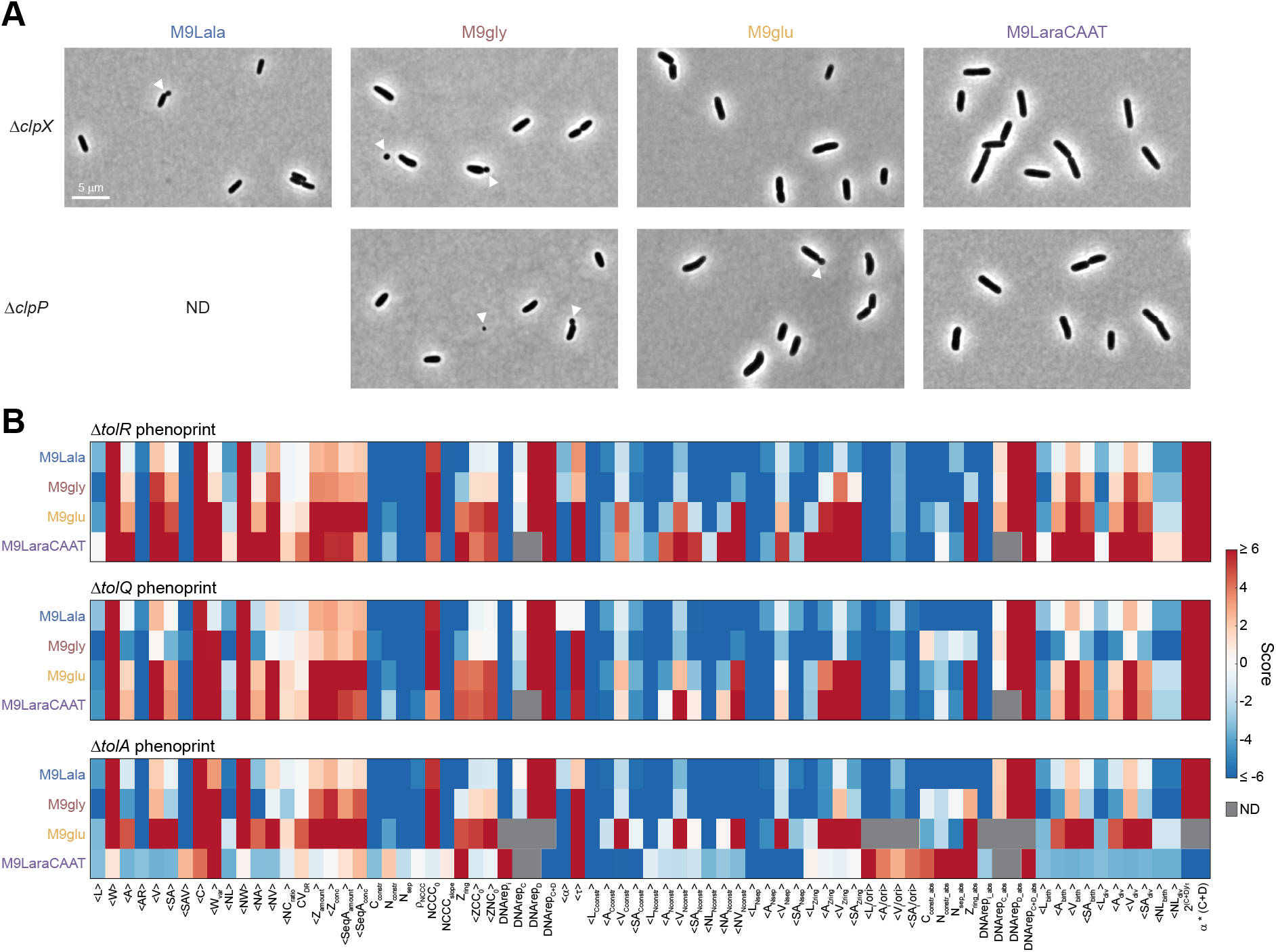
Examples of mutant phenotypes, related to Figure 7. A. Representative images of Δ*clpX* and Δ*clpP* strains in the indicated nutrient conditions. White arrows indicate minicells (or minicell formation). ND indicates that images and features of the Δ*clpP* strain could not be determined in M9Lala due to the lack of growth of the strain in this nutrient condition. B. Heatmap showing the normalized scores for each of the 77 extracted population level features of the indicated deletion strains. Grey boxes indicate DNA replication-related features that could not be determined (ND) because of the presence of more than two overlapping rounds of DNA replication.

