## Supplementary material for "Apparent simplicity and emergent robustness in bacterial cell cycle control": Key resources table

| REAGENT or RESOURCE | SOURCE | IDENTIFIER |
| --- | --- | --- |
| Bacterial and virus strains | | |
| *E. coli* MG1655 *seqA*::*seqA-mcherry ftsZ*::*ftsZ-venus^SW^* | Gray et al.^1^ | CJW6324 |
| Individual deletion mutants in *E. coli* MG1655 *seqA*::*seqA-mcherry ftsZ*::*ftsZ-venus^SW^* | This paper | NA |
| Chemicals, peptides, and recombinant proteins | | |
| Agarose | AmericanBio | Cat#AB00972-00500 |
| 4’,6-Diamidine-2′-phenylindole dihydrochloride (DAPI) fluorescent dye | Thermo Fisher Scientific | Cat#D1306 |
| Phusion high-fidelity polymerase | New England Biolabs | Cat#M0530S |
| LB | Fisher Scientific | Cat#BP1426-2 |
| BHI | Fisher Scientific | Cat#11469668 |
| NB | Fisher Scientific | Cat#10679125 |
| TSB | Fisher Scientific | Cat#10098983 |
| Oligonucleotides | | |
| Primers for strain verification/construction | This paper, Integrated DNA Technologies | See Table S2 |
| Software and algorithms | | |
| ImageJ | Collins^2^ | <https://imagej.nih.gov/ij/> |
| MATLAB | Mathworks | <https://www.mathworks.com/> |
| Oufti | Paintdakhi et al.^3^ | <https://oufti.org/> |
| Github code repository of the Jacobs-Wagner lab |  | <https://github.com/JacobsWagnerLab/published> |
