## Supplemental tables for "Apparent simplicity and emergent robustness in bacterial cell cycle control"

**Table S1. Growth medium composition, related to Figures 1-7 and STAR Methods.**

| **Abbreviation** | **Medium composition** |
| --- | --- |
| Brain Heart Infusion (BHI) broth | 200 g/l calf brain infusion, 250 g/l beef heart infusion, 10 g/l proteose peptone, 5 g/l NaCl, 2.5 g/l Na_2_HPO_4_, 2 g/l glucose |
| Lysogeny broth (LB) | 10 g/l NaCl, 5 g/l yeast extract, 10 g/l tryptone |
| M9 salts | 6 g/l Na_2_HPO_4_·7H_2_O, 3 g/l KH_2_PO_4_, 0.5 g/l NaCl, 1 g NH_4_Cl, 2 mM MgSO_4_, 0.1 mM CaCl_2_ |
| M9acet | 1x M9 salts, 0.2% acetate (sodium salt) |
| M9acetCAAT | 1x M9 salts, 0.2% acetate, 0.1% casamino acids, 1 µg/ml thiamine |
| M9αKG | 1x M9 salts, 0.2% α-ketoglutarate (sodium salt) |
| M9αKGCAAT | 1x M9 salts, 0.2% α-ketoglutarate, 0.1% casamino acids, 1 µg/ml thiamine |
| M9fuc | 1x M9 salts, 0.2% fucose |
| M9fucCAAT | 1x M9 salts, 0.2% fucose, 0.1% casamino acids, 1 µg/ml thiamine |
| M9fum | 1x M9 salts, 0.2% fumarate (disodium salt) |
| M9fumCAAT | 1x M9 salts, 0.2% fumarate, 0.1% casamino acids, 1 µg/ml thiamine |
| M9glcnac | 1x M9 salts, 0.2% N-acetylglucosamine |
| M9glcnacCAAT | 1x M9 salts, 0.2% N-acetylglucosamine, 0.1% casamino acids, 1 µg/ml thiamine |
| M9glu | 1x M9 salts, 0.2% glucose |
| M9gluCAAT | 1x M9 salts, 0.2% glucose, 0.1% casamino acids, 1 µg/ml thiamine |
| M9gly | 1x M9 salts, 0.2% glycerol |
| M9glyCAAT | 1x M9 salts, 0.2% glycerol, 0.1% casamino acids, 1 µg/ml thiamine |
| M9lact | 1x M9 salts, 0.2% lactose |
| M9lactCAAT | 1x M9 salts, 0.2% lactose, 0.1% casamino acids, 1 µg/ml thiamine |
| M9Lala | 1x M9 salts, 0.2% L-alanine |
| M9LalaCAAT | 1x M9 salts, 0.2% L-alanine, 0.1% casamino acids, 1 µg/ml thiamine |
| M9Lara | 1x M9 salts, 0.2% L-arabinose |
| M9LaraCAAT | 1x M9 salts, 0.2% L-arabinose, 0.1% casamino acids, 1 µg/ml thiamine |
| M9mala | 1x M9 salts, 0.2% malate (sodium salt) |
| M9malaCAAT | 1x M9 salts, 0.2% malate, 0.1% casamino acids, 1 µg/ml thiamine |
| M9malt | 1x M9 salts, 0.2% maltose |
| M9maltCAAT | 1x M9 salts, 0.2% maltose, 0.1% casamino acids, 1 µg/ml thiamine |
| M9mann | 1x M9 salts, 0.2% mannose |
| M9mannCAAT | 1x M9 salts, 0.2% mannose, 0.1% casamino acids, 1 µg/ml thiamine |
| M9pyr | 1x M9 salts, 0.2% pyruvate (sodium salt) |
| M9pyrCAAT | 1x M9 salts, 0.2% pyruvate, 0.1% casamino acids, 1 µg/ml thiamine |
| M9sorb | 1x M9 salts, 0.2% sorbitol |
| M9sorbCAAT | 1x M9 salts, 0.2% sorbitol, 0.1% casamino acids, 1 µg/ml thiamine |
| M9succ | 1x M9 salts, 0.2% succinate (disodium salt) |
| M9succCAAT | 1x M9 salts, 0.2% succinate, 0.1% casamino acids, 1 µg/ml thiamine |
| M9treh | 1x M9 salts, 0.2% trehalose |
| M9trehCAAT | 1x M9 salts, 0.2% trehalose, 0.1% casamino acids, 1 µg/ml thiamine |
| M9xyl | 1x M9 salts, 0.2% xylose |
| M9xylCAAT | 1x M9 salts, 0.2% xylose, 0.1% casamino acids, 1 µg/ml thiamine |
| Nutrient broth (NB) medium | 3 g/l beef extract, 5 g/l peptone |
| PYE | 2 g/l bacto-peptone, 1 g/l yeast extract, 1 mM MgSO_4_, 0.5 mM CaCl_2_ |
| Tryptic soy broth (TSB) | 17 g/l tryptone, 3 g/l soya-peptone, 5 g/l NaCl, 2.5 g/l K_2_HPO_4_, 2.5 g/l glucose |

**Table S2. Oligonucleotides used in this study, related to STAR Methods.**

| **Name** | **Sequence (5’ to 3’)** | **Use** |
| --- | --- | --- |
| SG105 | GTACTGATGGCTTCGGTCGTTC | Forward primer for checking *aceF* |
| SG106 | CTGGCGCTTGCTACCTTGC | Reverse primer for checking *aceF* |
| SG107 | ATCGAGTTGCTGCTTGGTGAGA | Forward primer for checking *rnlA* |
| SG108 | CACTCTGGACCGTTAGCGCAC | Reverse primer for checking *rnlA* |
| SG109 | GATATACTCAGCGGCAGGGAG | Forward primer for checking *hicB* |
| SG110 | CCTGACTTTCGAGCAACTGATAG | Reverse primer for checking *hicB* |
| SG111 | GATCGCGGTATTGTGGTGGTC | Forward primer for checking *pncA* |
| SG112 | GGTCGATCACGGTGTTTATGGT | Reverse primer for checking *pncA* |
| SG113 | GACTTCTCTCGCCGTCCAGC | Forward primer for checking *menB* |
| SG114 | GAAGGCATCTGCGGTAGCTC | Reverse primer for checking *menB* |
| SG115 | GACCTCCGCCTTTGATTTGAC | Forward primer for checking *ydaS* |
| SG116 | CCAACGGTGATAGATATTCTGCTG | Reverse primer for checking *ydaS* |
| SG117 | GTCCTGACATCATCCGCATC | Forward primer for checking *fucR* |
| SG118 | CAGGCACAGCTTGATGAACATG | Reverse primer for checking *fucR* |
| SG119 | GAATGTGTGAACGCTGCCGA | Forward primer for checking *ypfN* |
| SG120 | GTAAGCCTGGCGATGATGATGT | Reverse primer for checking *ypfN* |
| SG121 | CACGGCGACCTGGAAGAAG | Forward primer for checking *eutN* |
| SG122 | GCATTGCCACGCTTTTTAACC | Reverse primer for checking *eutN* |
| SG123 | GAAGATCAAAATATACGACCGCTC | Forward primer for checking *yibN* |
| SG124 | CACGATGGCAATACGGGCA | Reverse primer for checking *yibN* |
| SG125 | CGGATTGCTGCGAAAGAAAT | Forward primer for checking *waaQ* |
| SG126 | CGTGCGGCAACTGTTGATG | Reverse primer for checking *waaQ* |
| SG182 | GATGCGTTTCGTGGCGTTG | Forward primer for checking *yfdV* |
| SG183 | CATTATTTGTCATTTGCTTATGCCTT | Reverse primer for checking *yfdV* |
| SG184 | CACCGCAGAGATGTTACATTGATG | Forward primer for checking *yehE* |
| SG185 | GCTTTCTAAATGCAATTAAAATCAGGTT | Reverse primer for checking *yehE* |
| SG186 | CTCACCCCAGCCCTCACC | Forward primer for checking *entC* |
| SG187 | AGCGCGGTTTCACCAGGT | Reverse primer for checking *entC* |
| SG188 | TACTTACTCTCTTCAGCCTGACGG | Forward primer for checking *dapF* |
| SG189 | CATGTGCCACTCGACCAACG | Reverse primer for checking *dapF* |
| SG190 | GCTTTTCTGCCGTCGATTATG | Forward primer for checking *ybhR* |
| SG191 | CGTGGGATTATGCGCACTGTATT | Reverse primer for checking *ybhR* |
| SG192 | GAGCGTTTACCACCAGAGCAG | Forward primer for checking *tolA* |
| SG193 | AACACCAATAGGACGACCGGAA | Reverse primer for checking *tolA* |
| SG194 | ACCGCCATAGTATGAAACTGCC | Forward primer for checking *cysD* |
| SG195 | GTCCACCAGCAGAGCCAGAT | Reverse primer for checking *cysD* |
| SG196 | GATTGCTGGTCTGTTGCCGT | Forward primer for checking *atpE* |
| SG197 | GACAAACGCGATGGCCTG | Reverse primer for checking *atpE* |
| SG198 | GCGAAAGGGCTGGGTTATGA | Forward primer for checking *ybbD* |
| SG199 | GATATTCATCGTCTGAGCTATATGGC | Reverse primer for checking *ybbD* |
| SG200 | CTCTCATATGTTCGCCCGACT | Forward primer for checking *yfdQ* |
| SG201 | AAGCGGCAGATATTTGAAAGG | Reverse primer for checking *yfdQ* |
| SG202 | CGATCAGCGCCTGAAAATGC | Forward primer for checking *yjiL* |
| SG203 | GCAATGCAGCGCCAGACAC | Reverse primer for checking *yjiL* |
| SG204 | GAGAAAGCCGATGGGGTGA | Forward primer for checking *yacL* |
| SG205 | GATATGAAGGCGCTGTATGACA | Reverse primer for checking *yacL* |
| SG206 | CCGAAGTCTATAACAAGGATGGTAA | Forward primer for checking *yddK* |
| SG207 | GATGAATTGGGTGAAGTGCTGA | Reverse primer for checking *yddK* |
| SG208 | TTTACTTTGCGGCAGATGAACA | Forward primer for checking *pgm* |
| SG209 | GTGACACTATGACGACCAGACTCC | Reverse primer for checking *pgm* |
| SG217 | CCATTTCCGGCATCGACTCA | Forward primer for checking *sspA* |
| SG218 | ACGCGGAATGCCACCAAA | Reverse primer for checking *sspA* |
| SG241 | TAATGCGTCTTATCAGGCCTACAG | Forward primer for checking *iscR* |
| SG242 | CGAAACGGTGAGAACGGGAG | Reverse primer for checking *iscR* |
| SG243 | GATTTTGTCAGGCTTGCGGA | Forward primer for checking *stpA* |
| SG244 | TCTCAACCCTTGCCGTTAATG | Reverse primer for checking *stpA* |
| SG245 | CTCACGTGCTGCGAAATCATC | Forward primer for checking *hns* |
| SG246 | TCGTCATAACACCCTTGGCAC | Reverse primer for checking *hns* |
| SG247 | CGTAAATCAGGTAGTTGGCGTAAAC | Forward primer for checking *ihfA* |
| SG248 | CAGTTCGCCACGAGGAGTAAAC | Reverse primer for checking *ihfA* |
| SG249 | AAGGGTACCGAATTGCACGTAAA | Forward primer for checking *fis* |
| SG250 | AGTTCGGCAAGCGCATCAC | Reverse primer for checking *fis* |
| SG251 | ACGGTGATGACGATGAGGGA | Forward primer for checking *mraZ* |
| SG252 | GAGAAAGGTCCGTGGATGATG | Reverse primer for checking *mraZ* |
| SG253 | GAGTGGCGTCAGGCGATATAC | Forward primer for checking *ybaB* |
| SG254 | AGTGGCCGATTTCCGACAT | Reverse primer for checking *ybaB* |
| SG255 | CGTCTGGTATGCAGGTTGTGA | Forward primer for checking *hupB* |
| SG256 | AATAATCTTGAGCACGAGACTGTTT | Reverse primer for checking *hupB* |
| SG257 | TGCGGAAAAAGGAAGCGTAA | Forward primer for checking *minC* |
| SG258 | CGTTGCATCGCCCTGAAT | Reverse primer for checking *minC* |
| SG259 | GCAATCACAATCGGCGTCAG | Forward primer for checking *yfjW* |
| SG260 | GCTTTTCAACCGCTCAGTTCTG | Reverse primer for checking *yfjW* |
| SG261 | GCAACCGTTTTCACTTTCCGT | Forward primer for checking *opgC* |
| SG262 | AGTATGATGCGGCCTGGC | Reverse primer for checking *opgC* |
| SG263 | AATCACTGCCGGTTACATTGTAGT | Forward primer for checking *dksA* |
| SG264 | GGAACTTCACGAGGCGGGT | Reverse primer for checking *dksA* |
| SG265 | CTTCGCTGGGGAAACCTGTG | Forward primer for checking *wecC* |
| SG266 | CATGACACAGTCTGGCTGATGCT | Reverse primer for checking *wecC* |
| SG267 | TTTGTTGCAGACGACGAATTG | Forward primer for checking *mtn* |
| SG268 | ACGCCAGGCAATCACCAGAT | Reverse primer for checking *mtn* |
| SG269 | CCAAAGAATATGCTCTGCTGTCAC | Forward primer for checking *basS* |
| SG270 | TTGACCACGAGATTGCCGAT | Reverse primer for checking *basS* |
| SG271 | CTACTGGACGGGTGGCAACT | Forward primer for checking *hyaD* |
| SG272 | GATGGCTTCGCTCTGCTCAA | Reverse primer for checking *hyaD* |
| SG273 | CTTTCAAAATGACCGTTGCTCTCT | Forward primer for checking *yfdX* |
| SG274 | AAGAAGCGCCGGTGATTTACAA | Reverse primer for checking *yfdX* |
| SG275 | CGTCACGTAAAGCTTGTCGAGTT | Forward primer for checking *yfeC* |
| SG276 | AGGAGCGTATTCAGCAATTCTCTC | Reverse primer for checking *yfeC* |
| SG277 | CTCACGATAAGAAACGCATCCG | Forward primer for checking *ydcY* |
| SG278 | GGACACTTATACCTGGCTTGCTG | Reverse primer for checking *ydcY* |
| SG279 | GTAACGGTATCGAGATTGAACACG | Forward primer for checking *iscX* |
| SG280 | GCAGGTTGGGTAGAGAGGGTAATC | Reverse primer for checking *iscX* |
| SG281 | TCGTGAACCCTTACCGCCTG | Forward primer for checking *ycdX* |
| SG282 | ACGCATTGTAATCGGCAGAGA | Reverse primer for checking *ycdX* |
| SG283 | GCAGCCGTGGAGTAGCGAA | Forward primer for checking *ycbL* |
| SG284 | TGTGCGAAGCGATTGTGG | Reverse primer for checking *ycbL* |
| SG285 | GGTACATGGGGAAAGTGATAAAAATG | Forward primer for checking *wcaE* |
| SG286 | CGTAATCACCTAAGGTTAATTTCCAC | Reverse primer for checking *wcaE* |
| SG287 | CGACTTTATTCCCCTGGTATGTGT | Forward primer for checking *mdtE* |
| SG288 | CCATCAAGCCCATTCATATTTTG | Reverse primer for checking *mdtE* |
| SG299 | CTGCCGTTAGACCAGATGCTG | Forward primer for checking *hflC* |
| SG300 | ATCGGGCAAATTGGTCATCG | Reverse primer for checking *hflC* |
| SG301 | GATACATCGCCGTCCATTTACTC | Forward primer for checking *ydiV* |
| SG302 | AACAGCGACGATGGAATACTCAG | Reverse primer for checking *ydiV* |
| SG303 | ACATGGAATTGCGCCGCT | Forward primer for checking *yodC* |
| SG304 | GTCTCGTGCTGCACCCTGAC | Reverse primer for checking *yodC* |
| SG305 | GATTTGGTAGCTATGTCGCGTTT | Forward primer for checking *blc* |
| SG306 | CGATGATTGTCAGTATGGCGCTA | Reverse primer for checking *blc* |
| SG307 | CACCACCTGCGGAAGAAAATC | Forward primer for checking *yciS* |
| SG308 | CAGGTTTCCGAGCGTAAGGTG | Reverse primer for checking *yciS* |
| SG309 | CAAATTGAGCGCGATGCCTT | Foward primer for checking *yohF* |
| SG310 | GAAGCACTGGGTCTGGTTGATTC | Reverse primer for checking *yohF* |
| SG311 | GAGTCACGGCAGAGAAGAAACC | Forward primer for checking *ybgA* |
| SG312 | ACACTGGCGACAAAGTCATCC | Reverse primer for checking *ybgA* |
| SG313 | CACTGATATGAAGCCCGAACTCG | Forward primer for checking *fucP* |
| SG314 | CACACTCGACGGCAGCTCC | Reverse primer for checking *fucP* |
| SG315 | CCATATACAAGTAGTGCTGCGTAAA | Forward primer for checking *ileS* |
| SG316 | CCAGGATCACGCTAATACCAATC | Reverse primer for checking *ileS* |
| SG317 | GCAATCTTTTTGGACATAGTCGTG | Forward primer for checking *yhjH* |
| SG318 | CCACGGATAGCGAACGGAC | Reverse primer for checking *yhjH* |
| SG319 | GAGAAAATGACTTCCACGCCTTAC | Forward primer for checking *ytfA* |
| SG320 | CAATGGCGCTCTTACCGTGTC | Reverse primer for checking *ytfA* |
| SG322 | CGATCCCGCACTTTGTCAGG | Foward primer for checking *tolB* |
| SG323 | GTAAACGATGTTGTTCTGCTGCA | Reverse primer for checking *tolB* |
| SG324 | GAAACGTCGCGAGATTATTAAAGAA | Foward primer for checking *adeQ* |
| SG325 | ATCCGGACGATGCAAATAACG | Reverse primer for checking *adeQ* |
| SG326 | GACGATACGCTGAATTTCTGCA | Foward primer for checking *smf* |
| SG327 | GGATCCGAGGCCAGTTGC | Reverse primer for checking *smf* |
| SG328 | CAACCCTGGCGAACTGTTTG | Foward primer for checking *srlE* |
| SG329 | GATAGCGATGCTGCCCGAG | Reverse primer for checking *srlE* |
| SG330 | CCTACCGCGTATTATCTGACCA | Foward primer for checking *ybgO* |
| SG331 | GAGATGTACCGCGAGGAAACC | Reverse primer for checking *ybgO* |
| SG332 | GCTTCCCGCTGACCATTTCTG | Foward primer for checking *ybjI* |
| SG333 | GACCGCTTGTGGAACGCTGA | Reverse primer for checking *ybjI* |
| SG334 | CATCTTTTATGCTGGTTGGCTGT | Foward primer for checking *ydbL* |
| SG335 | AACTAATTGGCGTTGCATGTACTG | Reverse primer for checking *ydbL* |
| SG336 | CAGACAGCAGTAGCACCAAAGG | Foward primer for checking *rzpD* |
| SG337 | TCGCCAGACAAATTAACCCGTA | Reverse primer for checking *rzpD* |
| SG338 | ATGAAGAACGGCTGGCGCT | Foward primer for checking *ynjB* |
| SG339 | GCAGACGGGCGCACATAC | Reverse primer for checking *ynjB* |
| SG340 | GCAGTACGAGAAAGAAGCGCA | Foward primer for checking *glyS* |
| SG341 | CTTTTCTCTGTCTGCCTTTCGGT | Reverse primer for checking *glyS* |
| SG342 | GGACATAACCACGAGGAGCATG | Foward primer for checking *yfdY* |
| SG343 | CATCAGCGATGACGACCACAC | Reverse primer for checking *yfdY* |
| SG346 | CTGCGCGGCTCGACGATA | Foward primer for checking *tolQ* |
| SG347 | CGATCTTTCTCAACCACCACG | Reverse primer for checking *tolQ* |
| SG350 | ATCCAGCTTGGCCCATTCG | Foward primer for checking *dacA* |
| SG351 | GTGTAGTCACCTGGCGCATGG | Reverse primer for checking *dacA* |
| SG354 | CGTTTCAAGGCCAACCCGA | Foward primer for checking *tolA* |
| SG355 | CAACACCAATAGGACGACCGGA | Reverse primer for checking *tolA* |
| SG356 | GCAGCAAAGTCAGGCATTTATACTC | Foward primer for checking *yraH* |
| SG357 | CGTTTCTGCGCCTTTGTTTG | Reverse primer for checking *yraH* |
| SG358 | CGTCCGAGCGCCTGAGC | Foward primer for checking *glmM* |
| SG359 | GTTGCCGCGTACCAATGCTT | Reverse primer for checking *glmM* |
| SG368 | CTTTCTGTGCAGCAGCTTTGTCG | Foward primer for checking *yaaY* |
| SG369 | CAGCCGGGTCAGTCTTGCC | Reverse primer for checking *yaaY* |
| SG370 | CATAATGCGCGGTAGCTCACA | Foward primer for checking *ydcS* |
| SG371 | GCTCCCACGGTGGCAGATT | Reverse primer for checking *ydcS* |
| SG372 | CTGGATAAAGGCGCACTGGC | Foward primer for checking *rimK* |
| SG373 | GCAGATGCAGAGCCTGACAGTTATC | Reverse primer for checking *rimK* |
| SG374 | CAAAAGCGGCATGGGTATTGAC | Foward primer for checking *yfcU* |
| SG375 | TAACGGCGAGGTGATCTTGTTAC | Reverse primer for checking *yfcU* |
| SG376 | CCAGTGATGCGCAGTTATGTACC | Foward primer for checking *ypfH* |
| SG377 | CGATAAGTGGGACGATGACGAC | Reverse primer for checking *ypfH* |
| SG378 | ACGCAAAGCTTAACGGTCAGG | Foward primer for checking *fadR* |
| SG379 | GCAAAGAAGTCCTGAAGCATGTG | Reverse primer for checking *fadR* |
| SG380 | CTTCTTCCTGTCTCACGAAAATCC | Foward primer for checking *ygfZ* |
| SG381 | CAGCGTCGCATCAGGCAT | Reverse primer for checking *ygfZ* |
| SG382 | GTATTGATTGGCGTGCAGATTG | Foward primer for checking *srlB* |
| SG383 | CCACGCGACCAAAGATTTCA | Reverse primer for checking *srlB* |
| SG384 | CCTTGATATGCGTAGTCTGTGGCT | Foward primer for checking *yciU* |
| SG385 | GAATTACCGTCGCCGATCAAC | Reverse primer for checking *yciU* |
| SG386 | ATAGCCGCAAAGCCCTGGT | Foward primer for checking *ansB* |
| SG387 | AGCTTGAGAATGCCGTGATACTG | Reverse primer for checking *ansB* |
| SG388 | GCAAACTTCTCCAACAACGCA | Foward primer for checking *ihfB* |
| SG389 | TCCGCTAATTATGCATACACCGA | Reverse primer for checking *ihfB* |
| SG390 | GCTATTTCGACCGTTTGCAGAG | Foward primer for checking *ydhQ* |
| SG391 | ACCTTTACGGCTTACTGGGTGC | Reverse primer for checking *ydhQ* |
| SG392 | GTCCGGTACCCATTGTTATTGCT | Foward primer for checking *lsrG* |
| SG393 | GGAGACGAGTCGATGCCTGTTA | Reverse primer for checking *lsrG* |
| SG394 | ACTTCCCGTCAGGCGTTTGT | Foward primer for checking *wzzE* |
| SG395 | CTGTCCTGGCTGCATTATGTTG | Reverse primer for checking *wzzE* |
| SG396 | CGAGATCCTGCCACAGAATAGC | Foward primer for checking *elyC* |
| SG397 | CGATCAGTTACTGGCGATGGTC | Reverse primer for checking *elyC* |
| SG398 | GCACAAAGGCCCGTCACC | Foward primer for checking *clpP* |
| SG399 | GTCGCAGATATACACGGATGGAC | Reverse primer for checking *clpP* |
| SG400 | GTGGAATACGGTCTGGTCGATT | Foward primer for checking *clpX* |
| SG401 | AACCACCACATCGCGCAG | Reverse primer for checking *clpX* |
| SG402 | TGCCGCATCGTTTAGTTTTAGC | Foward primer for checking *yegP* |
| SG403 | CATAAACAGCATCTGCGCCA | Reverse primer for checking *yegP* |
| SG406 | TAATAGTTGCGCGGCGTGC | Foward primer for checking *ybgI* |
| SG407 | CTCCCACCAGCGTTGCAAAC | Reverse primer for checking *ybgI* |
| SG410 | CCTGCGTAGCAACCCGTCT | Foward primer for checking *ybgF* |
| SG411 | ACCAACTGAGCTAACGACCCAC | Reverse primer for checking *ybgF* |
| SG412 | CGCTGCGGTACACGACATCC | Foward primer for checking *pcm* |
| SG413 | GGTGCCGGTGGATTTGAAGTG | Reverse primer for checking *pcm* |
| SG414 | TCCCGGTGTTCAAGTGGCCT | Foward primer for checking *fur* |
| SG415 | TAAACGTGCTGAAGAAAGAGAAACAA | Reverse primer for checking *fur* |
| SG416 | TGCGTAATTATATGGGGCATCTG | Foward primer for checking *ybfG* |
| SG417 | GCACTCGACCGATAGTCAATGAG | Reverse primer for checking *ybfG* |
| SG418 | GGTCACTGGGTCCATGCTGAA | Foward primer for checking *ybfE* |
| SG419 | GCAACGTCTTTACCAAGCTGTTT | Reverse primer for checking *ybfE* |
| SG420 | CTTTAGCGTCACAGACATGAAATTG | Foward primer for checking *pdhR* |
| SG421 | ATTCGATCGCCTGGAGCC | Reverse primer for checking *pdhR* |
| SG424 | CAAGCCCTGGCATGATTGG | Foward primer for checking *waaR* |
| SG425 | CGTTACGCTTAACTTTCGGAGAA | Reverse primer for checking *waaR* |
| SG426 | GGATTCCTTGTATCAGTGCCGA | Foward primer for checking *waaO* |
| SG427 | CACCATCAAGATAATTAGCATCGAC | Reverse primer for checking *waaO* |
| SG428 | GCAACGAAGAACGCCTGGA | Foward primer for checking *rpoD* |
| SG429 | CATTGGTTCGACGCAGATTTG | Reverse primer for checking *rpoD* |
| SG430 | CTAACCCAGCGATCAAAAAAGC | Foward primer for checking *rplK* |
| SG431 | CACCTTGGGTAAATACGGCTACG | Reverse primer for checking *rplK* |
| SG432 | CAGATCTTCTTCAATACGCATGTGC | Foward primer for checking *coaE* |
| SG433 | CACGCCAGGCACGCCA | Reverse primer for checking *coaE* |
| SG434 | AGTTTTCGACTTCGGCACAGC | Foward primer for checking *yedE* |
| SG435 | GAATGGTCGGCCCGTCTT | Reverse primer for checking *yedE* |
| SG436 | CGTGAACGTGGCCCATGCT | Foward primer for checking *yehM* |
| SG437 | CTCAATCACCTGCTGCGGAAA | Reverse primer for checking *yehM* |
| SG438 | CCAGCGAAAAGCGTCATTATCC | Foward primer for checking *yehQ* |
| SG439 | GCTGAAGCATGGGGAATAATTGT | Reverse primer for checking *yehQ* |
| SG440 | GGTTAAGTAGCCAGCCCGAGG | Foward primer for checking *ydaF* |
| SG441 | TATCACTTCTCCTTGCCGTAACC | Reverse primer for checking *ydaF* |
| SG442 | GCAACCTTAGCCCGCTGG | Foward primer for checking *ecpB* |
| SG443 | CTGGCTACCGGCGAGATGAAT | Reverse primer for checking *ecpB* |
| SG446 | ACATGAACCAGATGGCGAATG | Foward primer for checking *eutJ* |
| SG447 | GCTGTCCGCAACTGCTCACC | Reverse primer for checking *eutJ* |
| SG448 | TCATGACTGTGCCGGTGCTG | Foward primer for checking *yfjU* |
| SG449 | GACTGCACTAACGACTGACGGCT | Reverse primer for checking *yfjU* |
| SG450 | ACGGATACTGGATTCGGGGAT | Foward primer for checking *yqeJ* |
| SG451 | GGATTTGCACCAGATGTTGTTGAT | Reverse primer for checking *yqeJ* |

**Table S3. Parameters for cell detection in Oufti, related to STAR Methods.**

| **Parameter** | **Value** |
| --- | --- |
| outCsvFormat | 0 |
| csvFileEdit | 0 |
| runSerial | 0 |
| maxWorkers | 12 |
| algorithm | subpixel |
| invertimage | 0 |
| interpoutline | 0 |
| interSigma | 0 |
| areaMin | 200 |
| areaMax | 5000 |
| splitregions | 1 |
| displayW | 0 |
| wShedNum | 5800 |
| cellwidth | 10 |
| wspringconst | 0.1 |
| rigidity | 0.8 |
| rigidityB | 4 |
| imageforce | 7 |
| attrCoeff | 0.4 |
| repCoeff | 0.6 |
| neighRep | 12 |
| attrPower | 14 |
| fitDisplay | 0 |
| fitMaxIter | 250 |
| moveall | 0.3 |
| fitStep | 0.3 |
| fitStepM | 0.6 |
| fitCondition | 0 |
| fitqualitymax | 0.85 |
| fsmooth | 40 |
| roiBorder | 22.5 |
| noCellBorder | 1 |
| meshStep | 1 |
| meshWidth | 14 |
| getmesh | 1 |
| splitThreshold | 0.4 |
| joindist | 5 |
| joinangle | 0.2 |
| joindilate | 1 |
| eqaldist | 1.5 |
| edgemode | 1 |
| erodeNum | 0 |
| openNum | 0 |
| invertimage | 0 |
| thresFactorM | 0.86254 |
| thresFactorF | 0.86254 |
| threshminlevel | 0.3 |
| edgeSigmaL | 0.5 |
| valleythresh1 | 0 |
| logthresh | 0 |

**Table S4. Parameters for nucleoid detection in Oufti, related to STAR Methods.**

| **Parameter** | **Value** |
| --- | --- |
| Manual background threshold | 0.2 |
| Background subtraction method | 3 |
| Background subtraction threshold | 0.1 |
| Background filter size | 8 |
| Smoothing range (pixels) | 3 |
| Magnitude of LOG filter | 0.1 |
| Sigma of PSF | 1.62 |
| Fraction of object in cell | 0.4 |
| Minimum object area | 50 |

**Table S5. Features considered in this study and their associated symbols, related to Figures 1, 7 and S7.**

| **Number** | **Feature (unit)** | **Symbol** |
| --- | --- | --- |
| 1 | Mean cell length (µm) | <L> |
| 2 | Mean cell width (µm) | <W> |
| 3 | Mean cell area (µm^2^) | <A> |
| 4 | Mean cell aspect ratio | <AR> |
| 5 | Mean cell volume (µm^3^) | <V> |
| 6 | Mean cell surface area (µm^2^) | <SA> |
| 7 | Mean cell surface-area-to-volume ratio (µm^-1^) | <SAV> |
| 8 | Mean cell circularity (µm) | <C> |
| 9 | Mean individual cell width variability | <W_var_> |
| 10 | Mean nucleoid length (µm) | <NL> |
| 11 | Mean nucleoid width (µm) | <NW> |
| 12 | Mean nucleoid area (µm^2^) | <NA> |
| 13 | Mean nucleoid volume (µm^3^) | <NV> |
| 14 | Mean nucleocytoplasmic ratio | <NC_ratio_> |
| 15 | Division ratio variability | CV_DR_ |
| 16 | Mean amount of FtsZ-Venus^SW^ signal (A.U.) | <Z_amount_> |
| 17 | Mean amount of FtsZ-Venus^SW^ concentration (A.U./µm^2^) | <Z_conc_> |
| 18 | Mean amount of SeqA-mCherry signal (A.U.) | <SeqA_amount_> |
| 19 | Mean amount of SeqA-mCherry concentration (A.U./µm^2^) | <SeqA_conc_> |
| 20 | Relative timing of cell constriction | C_constr_ |
| 21 | Relative timing of nucleoid constriction | N_constr_ |
| 22 | Relative timing of nucleoid separation | N_sep_ |
| 23 | Correlation (Spearman) between cell and nucleoid constriction | ρ_NCCC_ |
| 24 | Extent of nucleoid constriction at initiation of cell constriction (µm) | NCCC_0_ |
| 25 | Slope of relationship between cell and nucleoid constriciton | NCCC_slope_ |
| 26 | Relative timing of FtsZ ring formation | Z_ring_ |
| 27 | Mean FtsZ amount in newly-constricted cells (A.U.) | <ZCC_0_> |
| 28 | Mean FtsZ amount in cells with a newly-constricted nucleoid (A.U.) | <ZNC_0_> |
| 29 | Relative timing of DNA replication initiation (B period) | DNArep_i_ |
| 30 | Relative duration of DNA replication (C period) | DNArep_C_ |
| 31 | Relative time from DNA replication termination to cell division (D period) | DNArep_D_ |
| 32 | Relative duration of C + D period | DNArep_C+D_ |
| 33 | Maximum growth rate (h^-1^) | <α> |
| 34 | Minimal doubling time (h) | <τ> |
| 35 | Mean cell length at initiation of cell constriction (µm) | <L_Cconstr_> |
| 36 | Mean cell area at initiation of cell constriction (µm²) | <A_Cconstr_> |
| 37 | Mean cell volume at initiation of cell constriction (µm^3^) | <V_Cconstr_> |
| 38 | Mean cell surface area at initiation of cell constriction (µm²) | <SA_Cconstr_> |
| 39 | Mean cell length at initiation of nucleoid constriction (µm) | <L_Nconstr_> |
| 40 | Mean cell area at initiation of nucleoid constriction (µm²) | <A_Nconstr_> |
| 41 | Mean cell volume at initiation of nucleoid constriction (µm^3^) | <V_Nconstr_> |
| 42 | Mean cell surface area at initiation of nucleoid constriction (µm²) | <SA_Nconstr_> |
| 43 | Mean nucleoid length at initiation of nucleoid constriction (µm) | <NL_Nconstr_> |
| 44 | Mean nucleoid area at initiation of nucleoid constriction (µm²) | <NA_Nconstr_> |
| 45 | Mean nucleoid volume at initiation of nucleoid constriction (µm^3^) | <NV_Nconstr_> |
| 46 | Mean cell length at initiation of nucleoid separation (µm) | <L_Nsep_> |
| 47 | Mean cell area at initiation of nucleoid separation (µm²) | <A_Nsep_> |
| 48 | Mean cell volume at initiation of nucleiod separation (µm^3^) | <V_Nsep_> |
| 49 | Mean cell surface area at initiation of nucleoid separation (µm²) | <SA_Nsep_> |
| 50 | Mean cell length at FtsZ ring formation (µm) | <L_Zring_> |
| 51 | Mean cell area at FtsZ ring formation (µm²) | <A_Zring_> |
| 52 | Mean cell volume at FtsZ ring formation (µm^3^) | <V_Zring_> |
| 53 | Mean cell surface area FtsZ ring formation (µm²) | <SA_Zring_> |
| 54 | Mean DNA replication initiation length per *ori* (µm) | <L_i_/*ori*> |
| 55 | Mean DNA replication initiation area per *ori* (µm²) | <A_i_/*ori*> |
| 56 | Mean DNA replication initiation volume per *ori* (µm^3^) | <V_i_/*ori*> |
| 57 | Mean DNA replication initiation surface area per *ori* (µm²) | <SA_i_/*ori*> |
| 58 | Absolute timing of cell constriction (h) | C_constr_abs_ |
| 59 | Absolute timing of nucleoid constriction (h) | N_constr_abs_ |
| 60 | Absolute timing of nucleoid separation (h) | N_sep_abs_ |
| 61 | Absolute timing of FtsZ ring formation (h) | Z_ring_abs_ |
| 62 | Absolute timing of DNA replication initiation (B period) (h) | DNArep_i_abs_ |
| 63 | Absolute duration of DNA replication (C period) (h) | DNArep_C_abs_ |
| 64 | Absolute time from DNA replication termination to cell division (D period) (h) | DNArep_D_abs_ |
| 65 | Absolute duration of C + D period (h) | DNArep_C+D_abs_ |
| 66 | Mean cell length at birth (µm) | <L_birth_> |
| 67 | Mean cell area at birth (µm²) | <A_birth_> |
| 68 | Mean cell volume at birth (µm^3^) | <V_birth_> |
| 69 | Mean cell surface area at birth (µm²) | <SA_birth_> |
| 70 | Mean cell length at division (µm) | <L_div_> |
| 71 | Mean cell area at division (µm²) | <A_div_> |
| 72 | Mean cell volume at division (µm^3^) | <V_div_> |
| 73 | Mean cell surface area at division (µm²) | <SA_div_> |
| 74 | Mean nucleoid length at birth (µm) | <NL_birth_> |
| 75 | Mean nucleoid length at division (µm) | <NL_div_> |
| 76 | Average number of *ori* | 2^(C+D)/τ^ |
| 77 | Growth rate * DNArep_C+D_abs_ | α * (C+D) |
| **Additional features related to the cell cycle laws** | | |
|  | Mean cell length added between birth and cell constriction (µm) | < ΔL_constr_> |
|  | Mean time from initiation of cell constriction time to completion of cell division (h) | <T_constr_> |
